# Exploring the roles of memory replay in targeted memory reactivation and birdsong development: Insights from computational models of complementary learning systems

**DOI:** 10.1101/2024.10.28.620229

**Authors:** Lu Yihe, Tamás Földes, Penelope A Lewis

## Abstract

Replay facilitates memory consolidation in both biological and artificial systems. Using the complementary learning systems (CLS) framework, we study replay in both humans and birds through computational modelling. We investigate impacts of replay triggered by targeted memory reactivation during sleep and experiments examining how sleep affects the development of birdsong in young songbirds. We show that qualitatively realistic sleep effects can be captured by highly abstracted, idealised CLS models. Our modelling sheds theoretical insights on the mechanisms underlying both strengthening and weakening effects of targeted memory reactivation, and supports the empirical hypothesis that replay drives overnight performance deterioration and correlates positively with the final performance in birdsong development.

**Author summary:** Taking a computational approach, we investigated the roles of memory replay in two complementary learning systems (CLS) models capturing realistic sleep effects observed in two real-life experiments on targeted memory reactivation (TMR) and birdsong development respectively. Our two CLS models are abstract and identical in architecture, and they are distinct in terms of where replay is generated. While the TMR model produces replay samples using its hippocampus, the birdsong model does so using the sensorimotor cortex. We found that certain TMR effects could characterise different TMR models, which might account for individual differences in human subjects. The results of the birdsong model support the idea that the dramatic overnight oscillations in performance accuracy which are observed during birdsong development are mainly driven by memory replay, and that long-term performance gain can be achieved despite short-term performance deterioration during the early nights of development. As we studied the two experiments using the unified CLS framework, we discuss how replay contributes to sleep-dependent performance changes from the perspective of systems consolidation.

## 1 Introduction

Memory replay, reactivation and rehearsal can all refer to the virtual reenactment of an experience, possibly fragmented and disordered, after learning. While these terms may describe different phenomena, especially in the neuroscience literature, they are generally associated with memory consolidation in both biological and artificial systems [1–3].

Reactivation of past neural activity patterns by the central nervous system was first detected in hippocampus [4], and subsequently in neocortex [5] of the rat (mammalian) brain. Later studies [6, 7] identified hippocampal replay, i.e., reactivation of patterns following a consistent temporal order, and similar phenomena have been confirmed in the neocortex [8]. Depending on the regions, cortical replay can reflect either the same wakeful experience in coordination with hippocampal replay [8, 9], or an independent experience in parallel to it [10].

While an avian brain is anatomically distinct from the mammalian brain, both the hippocampal formation and another brain structure called hyperpallium are thought to perform similar roles to the mammalian hippocampus and neocortex, respectively [11–14]. In particular, replay has been identified in the brain of zebra finches, a species of songbirds, and its critical role in the song acquisition and development in the juvenile males have been experimentally verified [15, 16].

Functionally, hippocampal replay is thought to be crucial to memory consolidation [17, 18] (c.f., two broader reviews [19, 20]). After the initial formation of a new memory in the hippocampus, they are thought to continue to reverberate through the cortico-hippocampal loop for several iterations. Subsequently, the cortical representation of the memory is strengthened and stabilised, while the reliance on the hippocampus is reduced. Via replay, the hippocampus and the neocortex perform complementary roles in learning. The hippocampus is responsible for rapid acquisition of new experiences that are pattern-separated, while the neocortex can gradually build strong integrated memory representations.

Such replay-mediated cortico-hippocampal interactions inspired the establishment of the complementary learning systems (CLS) theory [21], which conceptualises the hippocampus as the fast learning and memory system, and the neocortex as the slow system. More specifically, the fast system is capable of rapid learning of specific episodic experiences or events, creating sparse, pattern-separated representations, allowing for the encoding of detailed memories, while the slow system is responsible for slowly integrating and compressing the knowledge acquired from the fast system into more structured, overlapping representations, extracting statistical regularities and forming generalised semantic knowledge. The fast system uses replay to transfer knowledge to the slow system. The CLS theory has inspired many artificial systems of learning and memory, especially artificial neural networks (ANNs), to implement replay-like algorithms [1], in order to mitigate catastrophic forgetting [22] (or the stability-plasticity dilemma [23]), that is, old memories can be completely erased when the memory system acquires new information.

While a growing body of empirical evidence from mammalian brains has identified the neural substrates and the behavioural effects of replay, supporting and modernising the CLS theory [24, 25], i.e., we still lack a mechanistic understanding of replay at a system level. We therefore set out to investigate two real-life examples, targeted memory reactivation (TMR) [26] and birdsong development [16] by simulating different, hypothetical replay schemes in two computational models, derived from the CLS theory, that are identical in architecture.

TMR is an experimental technique aiming to modulate memory replay and consolidation. In a typical TMR experiment, sensor cues (usually sounds or smells) are associated with wakeful memories and then re-presented during sleep in order to bias the brain towards replaying this specific memory. TMR has various effects on memory consolidation – in some cases strengthening memories, in other cases helping them to integrate together, or facilitating extraction of gist-type information from a corpus of memories. When applied in REM TMR can reduce the emotional arousal associated with memories. Due to these diverse impacts, this technique is seen as a promising tool for tackling clinical problems, where it could be useful to help in rebuilding memories after trauma (such as stroke) or in reducing the emotionality of upsetting memories in anxiety disorders and depression. However, application of TMR currently requires experimenters to stay awake most of the night monitoring the sleep of participants, and as such it is highly labour-intensive, and impractical as a multi-night manipulation. We are therefore motivated to use computational modelling to bypass these practical challenges to gain a better understanding of the mechanisms of TMR, and in particular to ask how replay affects learning performance beyond mere strengthening of memories as observed in [27]. By shedding light on the optimisation of TMR, we would potentially increase its values in clinical applications.

That replay is not merely about memory strengthening is also corroborated by the observations from the birdsong development experiment [16]. We find this experiment on how juvenile zebra finch males learn songs by imitation particularly intriguing, because, while the experiment confirmed the positive correlation between their final performance and the total amount of replay throughout their development, on a daily basis their performance deteriorated overnight after sleep, presumably due to replay. This seemingly paradoxical phenomenon cannot be explain with mere repetition by veridical hippocampal replay, and, based our simulation, we argue non-veridical, generative replay provides extra information (noise) for the system to consolidate, which brings an adverse effect on the performance in the short term, but benefits learning (by making the system more robust) in the long term.

On top of addressing different research questions in the two real-life examples, we aim to demonstrate the generic effects of replay on learning and memory, which is one of our motivations to use the same architectures for the two CLS models for TMR and birdsong development. In both models the fast system is considered to be embodied by a *stochastic simple memory* (SSM) [28, 29], and the slow system by a *restricted Boltzmann machine* (RBM) [30, 31] (Fig 1). These two ANNs were chosen based primarily on their functional suitability (Section 4.1.3 and 4.1.2; similar choices were made in [17]). The SSM rapidly stores individual training samples as episodic memories, and the RBM gradually learns category prototypes from large number of samples. Their abstract, minimal neural architectures are appropriate given the similarities between mammalian and avian brains at the system level, allowing us to overlook the anatomical differences and to focus on different replay schemes.

**Fig 1.**
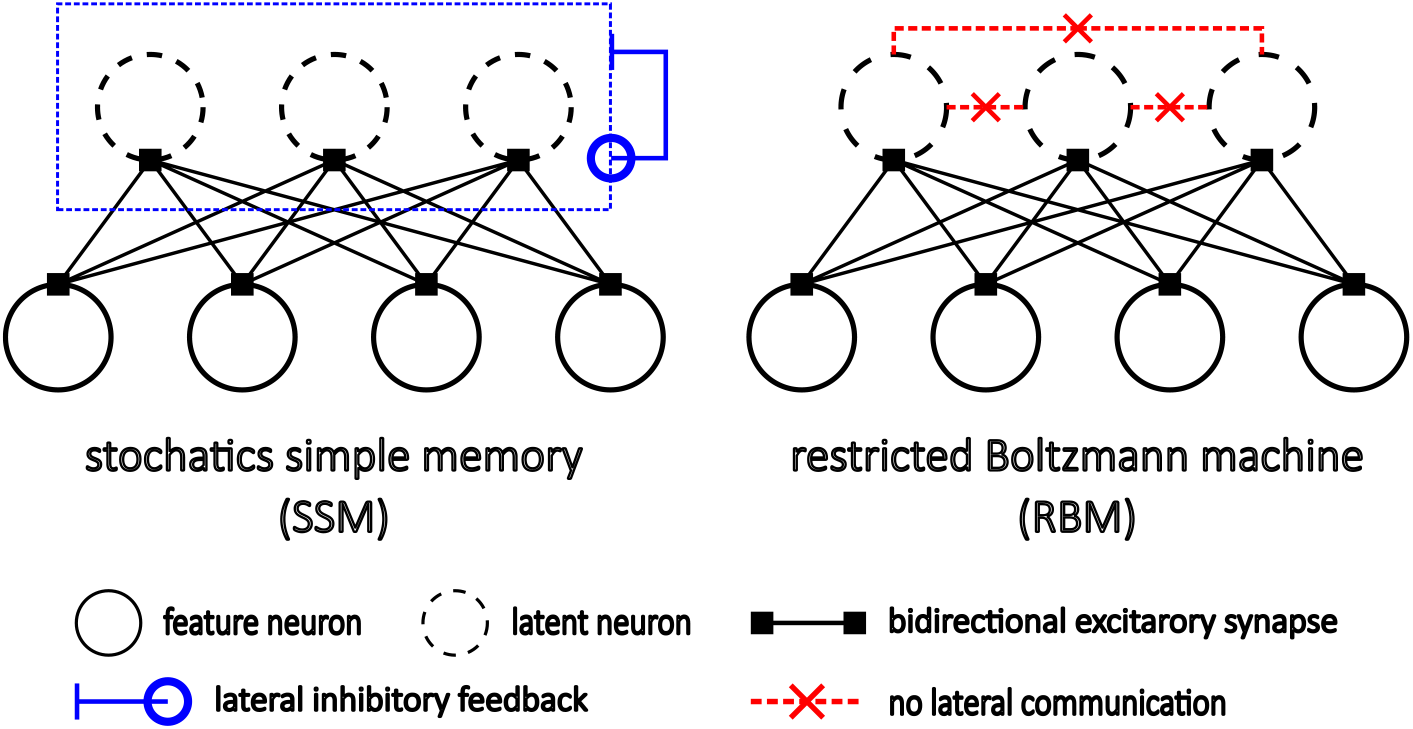
Schematics of the stochastic simple memory (SSM) and the restricted Boltzmann machine (RBM). Both neural networks consist of two layers of artificial neurons, a feature layer exchanging information with environment, and a latent layer communicating only with the feature layer via symmetric synapses. There are no lateral synapses between neurons within either layer. However, the latent neurons of the SSM activate competitively in a noisy, winner-take-all manner. Obeying their own learning rules, these two networks can acquire memories when training data are presented to their feature layer. They can produce replay samples as if performing memory retrieval, and consolidate replay samples as in training. Technical details of the models are summarised in Section 4.1.

Our two CLS models differ in terms of what information the fast system’s (hippocampal) replay contains. Our first CLS model for the TMR experiment (Sections 2.1.1 to 2.1.3) subscribes mainly to the classical CLS theory, as replay samples are produced by its hippocampus, and then transferred to the neocortex for consolidation. In contrast, when studying the birdsong development experiment (Sections 2.2.1 to 2.2.3), we test an alternative hypothesis that the fast system is responsible for indexing memory and its replay samples are effectively indices that initiate replay in the slow system.

## 2 Results

### 2.1 Targeted memory reactivation (TMR)

Real-life TMR experiments are mostly the same as a typical sleep experiment, in which participants are instructed to learn to perform a cognitive task and then tested before and after sleep, except that some sensory cues not directly relevant to the task are additionally presented to the participants during the wakeful training session, prompting them to form associations between the cues and certain ‘targeted’ task materials (determined by the experimenter), and that the cues are re-presented during the sleep period. The technique is called ‘targeted memory reactivation’, as it leads to excessive memory replay and enhanced task performance regarding the targeted task materials [26, 32–36].

We aimed to qualitatively reproduce such results in a simulated TMR experiment using an idealised complementary learning systems (CLS) model (Fig 2A). Specifically, the paired-associate learning task is considered, which gives participants multiple pairs of items, (*A, B*), to memorise, and subsequently tests how well they can recall a specific item, *A*, given its paired item, *B*. Instead of specifying the explicit modality of such items, given the abstract nature of our models, the items were assumed to be neurally encoded as a set of independent features, permitting a mathematical representation as a binary vector, and, similarly, any extra sensory cue was encoded in a vector, *C*. Consequently, any sample presented to our model was considered to be a longer vector concatenating the three parts. Explicitly, every training sample took the form of [*A*|*B*|*C*], and was randomly instantiated from one of the category prototypes of the synthetic data (Section 4.2.1). When testing the model, we probed the model by presenting [?|*B*|?], where ? denotes fully randomised encoding, as *B* was presented but *A* and *C* were not.

**Fig 2.**
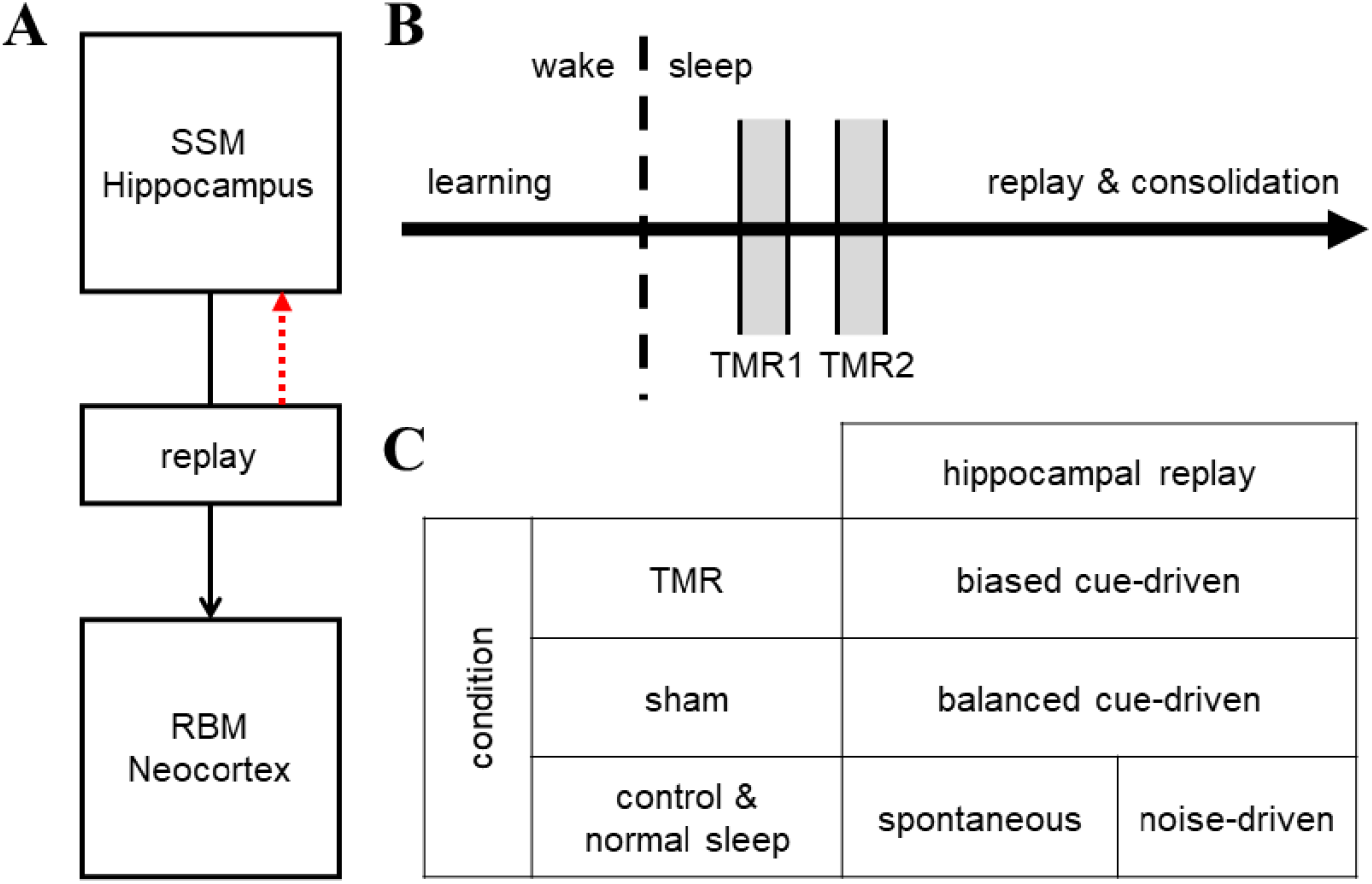
Model architecture and experimental protocol for targeted memory reactivation (TMR). (**A**) The idealised CLS model is composed of a hippocampus modelled by an SSM and a neocortex by an RBM. The red dashed arrow denotes the possibility that the hippocampus may be capable of memory consolidation via its own replay. (**B**) The model acquires new training samples, i.e., item pairs, coupled with sensory cues only in the wake session, and in sleep it performs memory replay and consolidation. There are two brief periods in sleep when TMR cues are presented. (**C**) 2 replay schemes in normal sleep, *spontaneous* and *noise-driven*, and 2 plasticity schemes, *plastic* and *stable*, are considered, yielding 4 models in total. TMR cues, similar to sensory cues in training, change the model’s replay mechanism from a normal mode to a cue-driven mode. Under the TMR condition, all the cues are associated with a particular item pair. Under the sham condition, the cue distribution is uniform with respect to all the pairs.

More specifically, the hippocampus was assumed to store training samples in wake, and to replay them in sleep, and the neocortex to start memory consolidation via hippocampal replay in sleep (Fig 2B). As all the memories was expected to eventually transfer from the hippocampus to the neocortex, only the neocortex was probed in tests, and we determined the model performance by measuring how similar to *A* the first part of its response was (Section 4.2.2).

We examined in total four different models in a two-dimensional design space, concerning two replay schemes and two plasticity schemes. The hippocampal replay was either *spontaneous* or *noise-driven*. If spontaneous, the hippocampus would choose randomly episodic memories as replay samples. If noise-driven, the hippocampus would produce replay samples by its cued-recall mechanism, using some random noise as the cue signal. In addition, the hippocampal memory could be *plastic* or *stable*, depending on whether or not the hippocampus performed memory consolidation via its own replay.

We note the exact neural substrates of these replay schemes are open to interpretations and not of our primary concern. What matters is that they are plausible in real brains. Nevertheless, we highlight that the four models are idealised in a mutually exclusive manner, whereas real brains are likely to fall on a spectrum charaterised by these idealisations, especially considering individual differences and noise at all levels.

In addition, in all the models, the hippocampal memory was constantly erode by a small amount of memory downscaling (Section 4.1.5).

#### 2.1.1 TMR manipulation varies paired-associate performance

We emulated the TMR manipulation by forcing the models to switch its hippocampal replay from the normal mode (i.e., spontaneous or noise-driven) to the cue-driven mode, which produced replay samples by the hippocampal cued-recall mechanism. Under the TMR condition, we re-presented only the sensory cues that had been coupled with the targeted item pair during 2 brief periods to the models, exactly as if it was a real-life TMR experiment. We also re-presented sensory cues under the sham condition, but uniformly for all the pairs. Under the control condition, there were no cues re-presented, and thus the hippocampal replay did not switch between the modes. The specific replay mechanisms under all the 3 conditions are summarised in Fig 2C.

As expected, the TMR manipulation introduced significant performance differences between the targeted memory and the non-targeted ones (Fig 3). Specifically, the performance on the targeted memory was significantly better than the non-targeted ones only under the TMR condition, and this difference persisted in time except in the plastic and spontaneous model (Fig 15). The effect, as observed in our simulations, was overwhelmingly significant compared to what has been observed in real-life TMR experiments primarily for two reasons:

**Fig 3.**
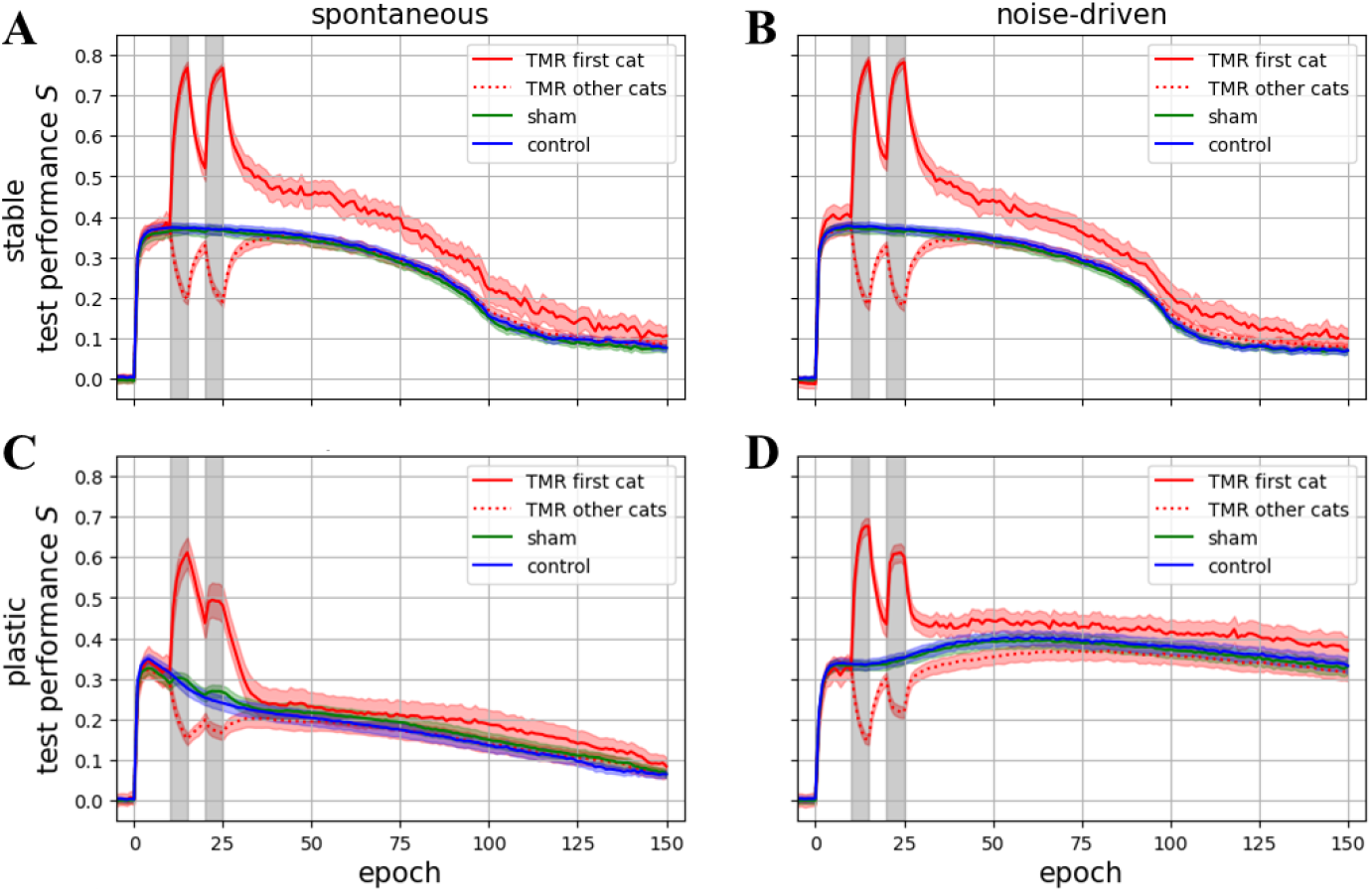
Test performance varied by TMR in 4 different models. A model’s hippocampus can be either stable (**AB**) or plastic (**CD**) in sleep (Fig 2A), and its replay can be either spontaneous (**AC**) or noise-driven (**BD**) (Fig 2C). Mean and 95% confidence interval from 100 trials are depicted.

- We gathered data from a relatively large number of simulated trials (100 repetitions) for each model. While there was noise in the simulations, each model was nearly identical across all the trials.
- The execution of replay and consolidation in all the models were perfect. The models were assumed to produce a constant number of replay samples per epoch, and consolidated all of them. In addition, they could switch to its cue-driven mode and back to the normal mode whenever necessary.

From now on we will focus on the analysis of the TMR effects, which is defined to be the performance differences between the TMR and sham conditions and the overall performance under the control condition. We are specifically interested in the post-cueing TMR effects, as it is trivial to observe strong TMR effects during the cueing periods.

Lastly, we note that the behaviours of the two models with the stable hippocampus are nearly identical, and that the performance kept decaying, and dropped to a relatively low level (*S <* 0.1; still significantly higher than the chance level *S* ≈ 0) after around the 110-th epoch in all the 4 models, except the much greater performance of the plastic and noise-driven model (Fig 3D). We discuss these 2 points in Sections 3.3 and 3.4.

#### 2.1.2 Different models manifest characteristic post-cueing TMR effects

Before analysing the TMR effects depicted in Fig 3 in more detail, we note that there was no strong evidence that the cueing manipulation (under the TMR or the sham condition) led to any long-lasting effect on the overall performance (Fig 16).This could be due to the brevity of the cueing periods, which introduced a relatively small number of biased samples among all the replay samples throughout sleep. Nevertheless, the TMR effects on the performance on the targeted pair in all the 4 models are persistently large and significant (Cohen’s *d >* 1 and *p*-value *<* 10^−50^ at nearly all the sleep epochs in Fig 4).

**Fig 4.**
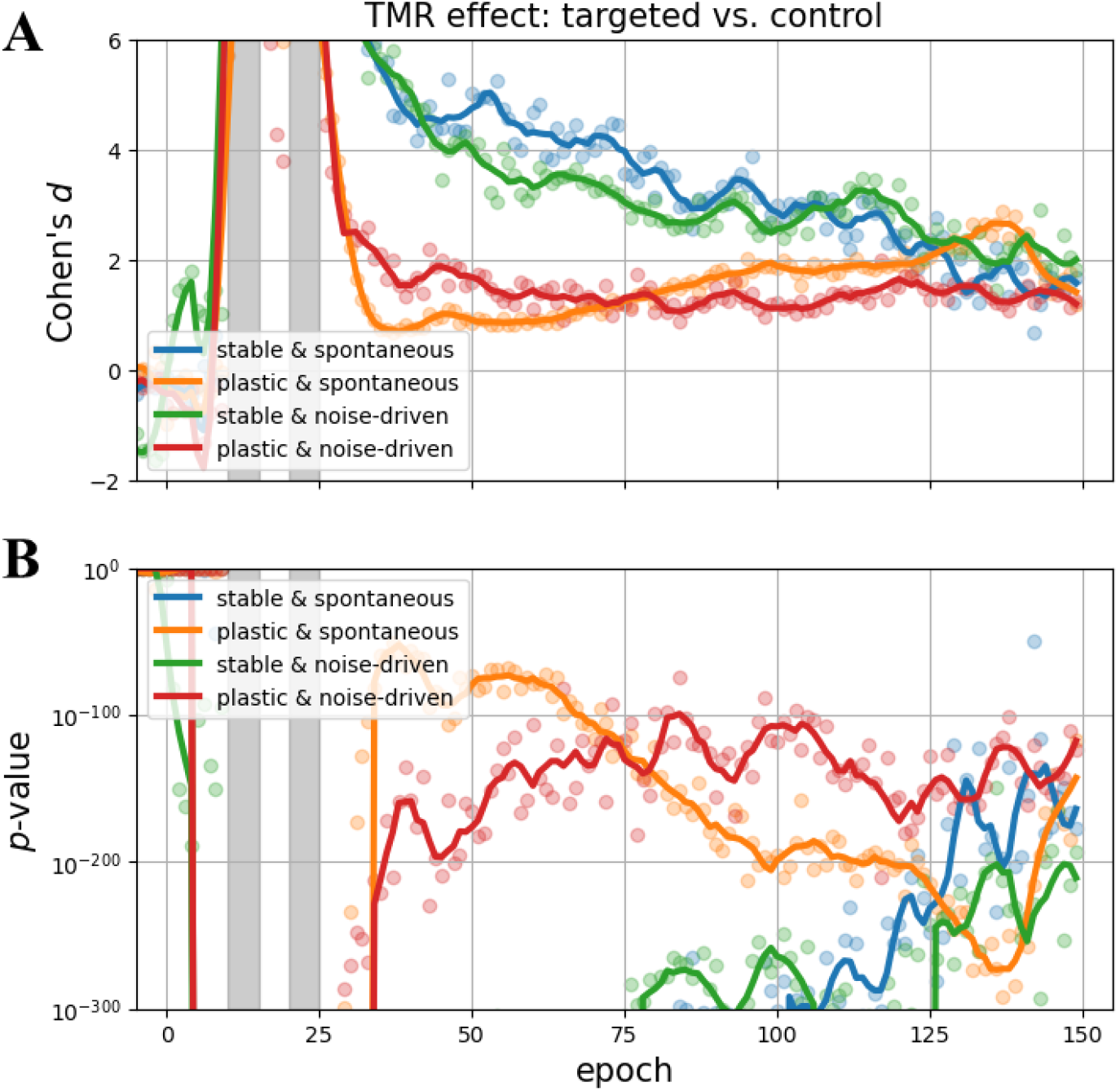
TMR effect on targeted memory under TMR condition. Both Cohen’s *d* (**A**) and *p*-value from a one-sided *t*-test (**B**) were computed with the data across all the epochs shown in Fig 3 (transparent dots), and then smoothed (solid lines). The alternative hypothesis of the *t*-test was that the performance on the targeted memory was better than the overall performance under the control condition.

Intriguingly, while the effect size decayed throughout sleep in the 2 stable models, it stayed at approximately the same level across sleep in the plastic and noise-driven model, and expressed a growing trend in the plastic and spontaneous model (Fig 4A). That model is thus more compatible than the others with the empirical observation that the TMR effect peaked after more than one week [37, 38], rather than the subsequent morning.

Interestingly, only the plastic and noise-driven model expresses the behaviour that the performance gain in the targeted pair was obtained at the cost of the memory loss in the non-targeted pairs (Fig 5), which is consistent with most empirical observations. We highlight that memory loss is not necessarily unfavourable, as a memory can be positive or negative. For example, the TMR manipulation has been shown to help patients reduce traumatic memory [39]. Thus, the plastic and noise-driven model might be a more suitable model for such patients than the other models.

**Fig 5.**
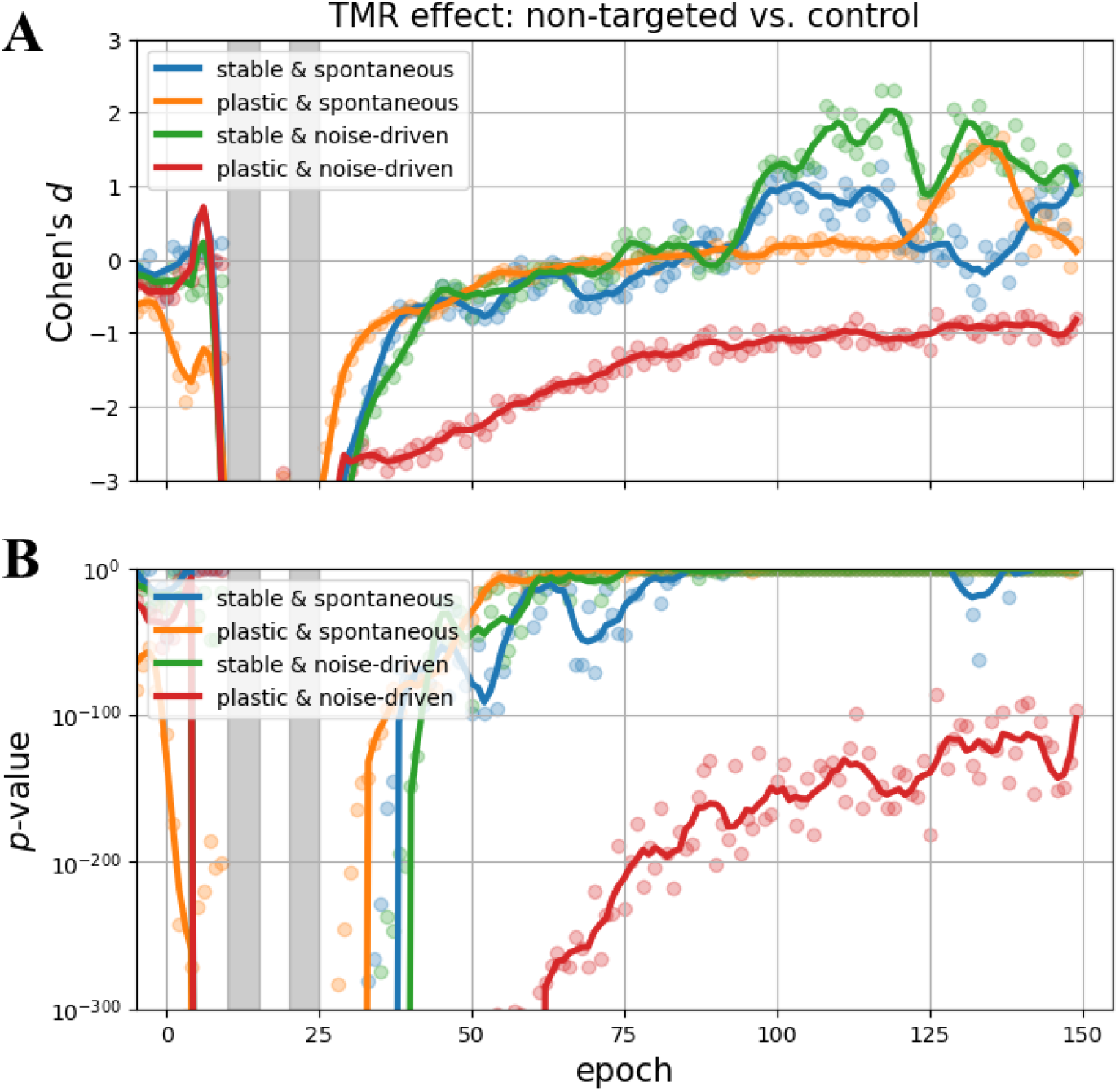
TMR effect on non-targeted memory under TMR condition. Both Cohen’s *d* (**A**) and *p*-value from a one-sided *t*-test (**B**) were computed with the data across all the epochs shown in Fig 3 (transparent dots), and then smoothed (solid lines). The alternative hypothesis of the *t*-test was that the performance on the non-targeted memory was worse than the overall performance under the control condition.

We note that the other models seemed to manifest significant performance gain in the late epochs, but consider this to be a statistical artifact, because the possible range for the variance of performance was largely reduced while the mean decayed close to chance level.

For completeness, we checked the TMR effect under the sham condition. Only the plastic and spontaneous model showed a large performance gain, whereas the sham manipulation was slightly detrimental in all the other models (ignoring the effects in the last epochs for the same reason as above; Fig 6).

**Fig 6.**
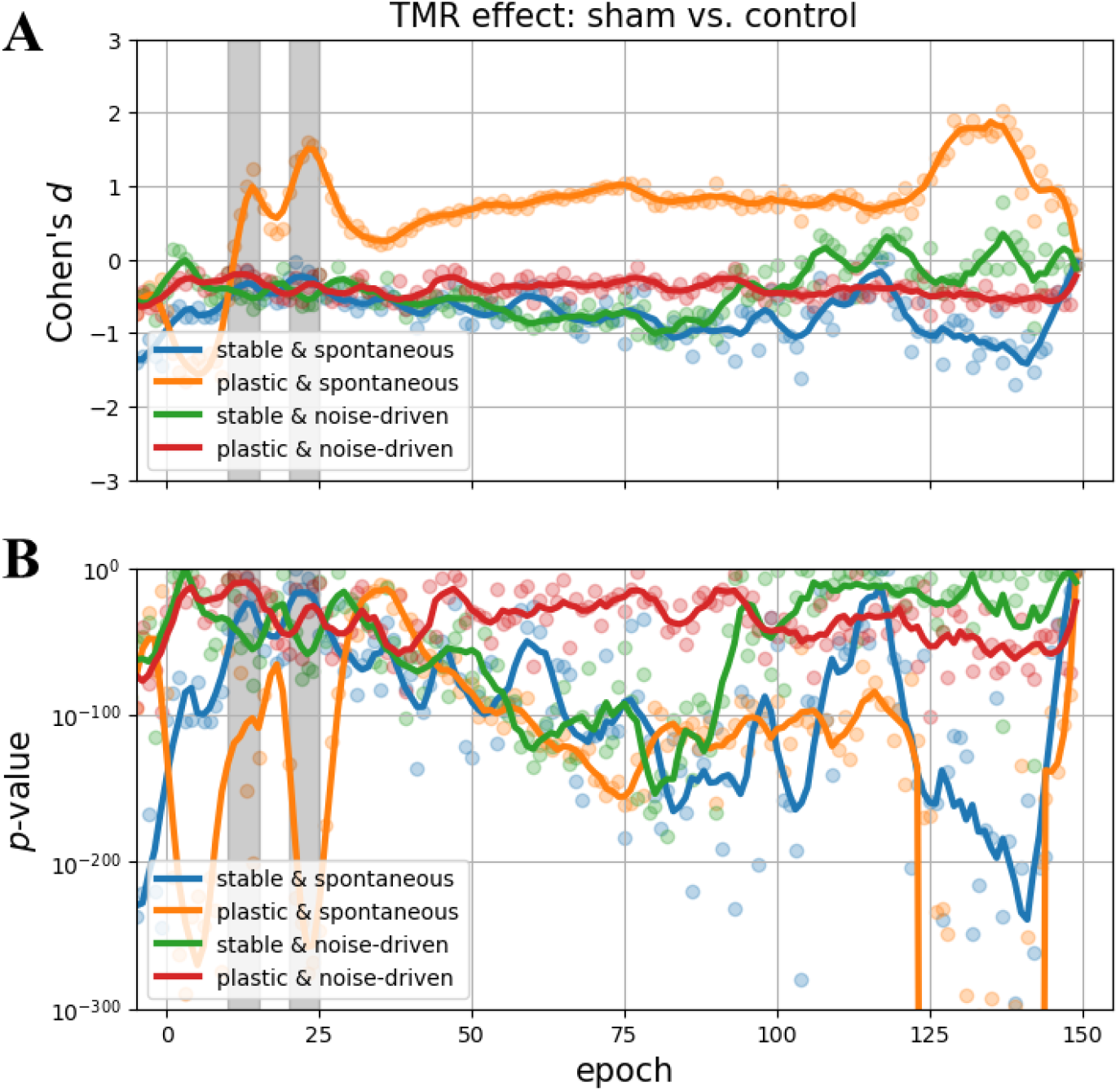
TMR effect under sham condition. Both Cohen’s *d* (**A**) and *p*-value from a two-sided *t*-test (**B**) were computed with the data across all the epochs shown in Fig 3 (transparent dots), and then smoothed (solid lines). The alternative hypothesis of the *t*-test was that the performance on the overall performance under the sham and the control conditions were different.

The TMR effects on all the models are summarised in Table 1. While the 2 stable models manifested qualitatively similar behaviours, the 2 plastic models were different from them and from each other.

**Table 1.** Post-cueing TMR effects on test performance.

| model | targeted | non-targeted | sham |
| --- | --- | --- | --- |
| stable & spontaneous | + ↘ | 0 | — |
| stable & noise-driven | + ↘ | 0 | — |
| plastic & spontaneous | + ↗ | 0 | + |
| plastic & noise-driven | + → | — | — |

The signs, + and —, denote statistically significant, positive and negative TMR effects, respectively, and 0 insignificant effects. The arrows indicate the main trends of the effect sizes in magnitude varying in time.

Although a real brain is likely to be characterised by a stochastic mixture of all the features in our modelling design space, it can be inferred from the models’ behavioural differences that:

- While a TMR manipulation permits performance gain in the targeted memory, this effect is only long-lasting, or even growing, if the hippocampus consolidates its own replay.
- A TMR manipulation can be effective at reducing non-targeted memory, if the hippocampus is relatively susceptible to external interference.

#### 2.1.3 TMR introduces weak discrepancy in replay

The previous sections have shown that TMR manipulations were able to vary performance of our models as in real-life experiments, and that the different replay schemes were ultimately responsible for the distinct TMR effects manifested by the models. Here we attempt to answer a more subtle question:

- How did the TMR manipulations, via changing replay, modulate the model memory and consequently the performance?

We measured replay samples in frequency and exactness (Section 4.2.3), and focussed on the differences between the targeted memory under the TMR condition and the average memory under the control condition (Fig 7), as the measures for the non-targeted memories and the sham condition are not significantly different from those under the control condition (Fig 17 and 18).

**Fig 7.**
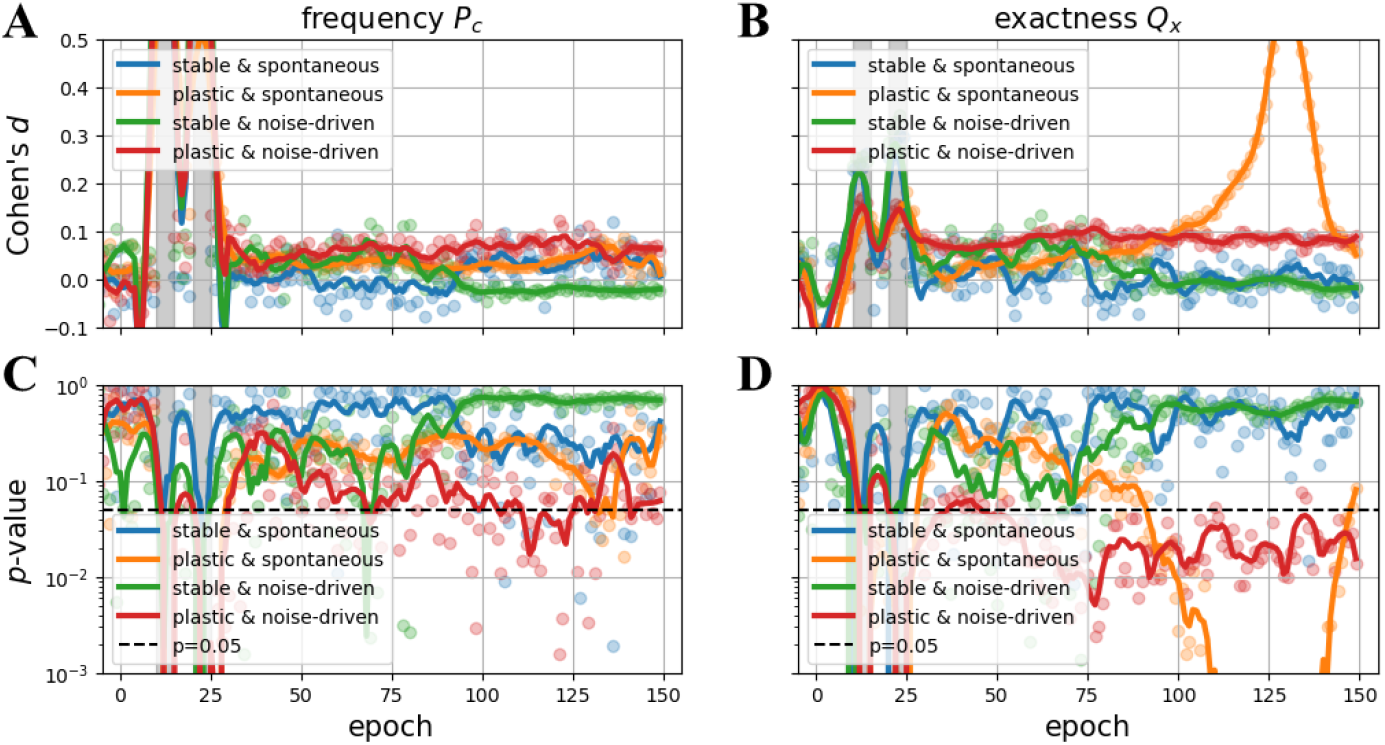
TMR effect on replay of targeted memory. Both Cohen’s *d* (**AB**) and *p*-value from a two-sided *t*-test (**CD**) were computed with the data of replay frequency *P*_*c*_ and replay exactness *Q*_*x*_ across all the epochs shown in Fig 17 and 18 (transparent dots), and then smoothed (solid lines). The alternative hypothesis of the *t*-test was that the replay measures of the targeted memory was different from those under the control condition.

It is clear from Fig 7BD that the replay exactness of the targeted memory was only significantly improved in the plastic and noise-driven model,, despite a small effect size, while there might have been an insignificant tendency to a minor gain in replay frequency (Fig 7C). There were no significant gains in either measure for the other models, except for the last epochs of the plastic and spontaneous model, when its memories were completely obliterated under the control condition (a phenomena also observed for other memories; Fig 18). Similar to our discussion in Section 2.1.2, the diminishing variance of replay exactness, rather than changes to the mean, was the dominant factor for this exceptional late-epoch effect.

Nevertheless, all the models manifested strong TMR effects on their test performance (Fig 4), which implies that:

- Either, the 2 replay measures are not informative enough, e.g., a comprehensive measure combining frequency and exactness is necessary.
- Or, the small replay bias introduced in the brief cueing periods were sufficient for the neocortex to form memory biased towards the targeted pair.

While being unable to exclude the first possibility (as we intentionally chose the 2 simple measures; Section 4.2.3), with hindsight we consider the second explanation more plausible (as discussed in Section 3.1). Specifically, the RBM in our models was sensitive to bias in training samples, especially in its early learning epochs, which probably accounts for the significant TMR effects.

As the only reason for us to choose the RBM as a proxy for the idealised neocortex was its statistical, slow learning property, we should be cautious about interpreting the TMR effects observed from the models. A different neocortical model may have manifested quantitatively different results.

Qualitatively, we consider the 2 plastic models’ capacity for maintaining (or increasing) the gain in replay exactness the explanation for them to manifest a long-lasting (or growing) TMR effect. While the initial replay bias may have been large enough to modulate the RBM’s performance, the replay samples were mostly balanced after cueing, and thus the TMR effects in the 2 stable models kept decaying. In contrast, when the hippocampus was plastic, the initial bias was weakly preserved throughout the following sleep epochs. The neocortex could thus consolidate more accurate replay samples of the targeted memory, leading to a better performance.

### 2.2 Birdsong development

Juvenile male zebra finches develop their song by imitating a template song, typically composed of a handful syllables. By measuring the similarity between such songs and templates in an experimental environment, experimenters [16] found that:

1. The songbirds sang less accurately after a night of sleep than on the previous evening, but rapidly recovered their performance and went on to improve more through further practice in daytime.
2. This overnight performance deterioration was largest in the early days of learning, and gradually diminished throughout development.
3. Remarkably, the extent of overall overnight deterioration was positively correlated to final developmental at the end of the developmental period.^1^

In this section we will focus on this performance oscillation, e.g., improvement during wake and deterioation overnight, in birdsong development. We simulated this phenomenon using an idealised CLS model consisting of a hippocampus and a sensorimotor cortex (Fig 8A) across 50 day-night cycles. The model was provided with training samples in wake, and it performed memory consolidation via replay in sleep.

**Fig 8.**
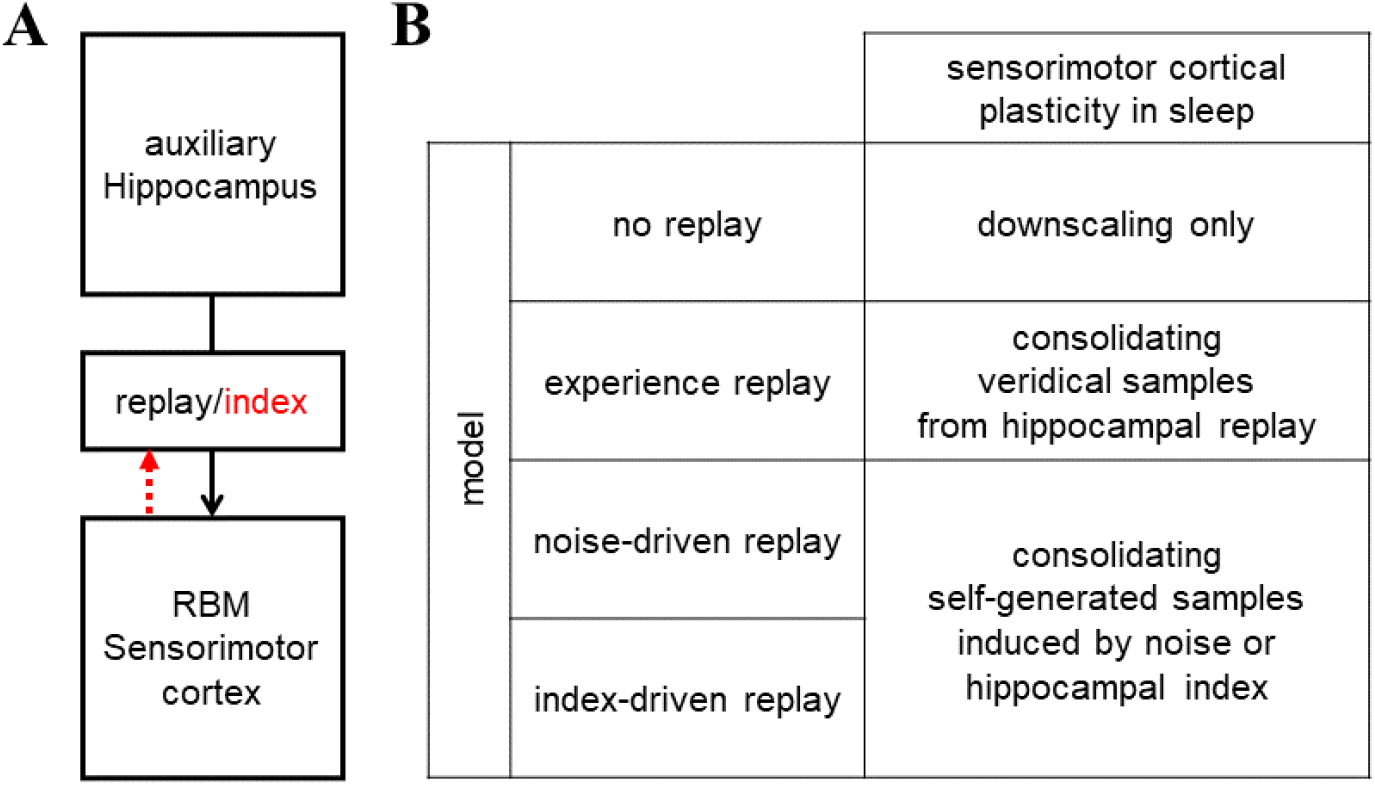
Model architecture and replay mechanisms for birdsong development. (**A**) The idealised CLS model is composed of a hippocampus modelled by an auxiliary storage and a sensorimotor cortex by an RBM. The red dashed arrow denotes the possibility that the sensorimotor cortex may generate its own replay induced by the reactivation of hippocampal index. (**B**) The sensorimotor cortex learned new samples in wake, and consolidated replay samples in sleep. Memory downscaling was implemented in all the models.

Specifically, each training sample, encoding a syllable, was assumed to be a paired pattern of auditory, *A*, and premotor signals, *C*, and a batch of them with independent auditory features and orthogonal sensorimotor activation was thus considered a representation of a song (Section 4.2.1). The assumption of orthogonality was made in light of the sequential neural activation for different syllables (and general motor signals) generally observed in hippocampus (not specific to singing) [6, 7, 15, 40–42]. Similar to our simulated TMR experiment, the CLS model was trained with synthetic data in the form of [*A*|*C*] vectors, and subsequently tested with [?|*C*], where *C* denotes the correct encoding and ? the fully randomised.

The CLS model was similar to the previous one for studying TMR (Fig 2A), with the hippocampus capable of veridical memory acquisition and replay, and the sensorimotor cortex also modelled by an RBM. Specifically, the sensorimotor cortex was tasked to learn the auditory-motor pairing as the neocortex in the previous sections to learn the association between task-relevant items and sensory cues.

Despite the architectural similarities with the TMR models, this CLS model for birdsong development was considered to produce replay samples from its sensorimotor cortex (rather than the hippocampus). The hippocampus was instead responsible for encoding and reactivating orthogonal premotor signals that were indices for the learned syllables, rather than the full representation [41, 43], and thus we named such replay *index-driven*. For comparison, we also considered *noise-driven* replay with such index completely randomised, and two other replay schemes with no sensorimotor cortical replay (Fig 8B). We note that *experience replay* here is by definition identical to spontaneous (hippocampal) replay in previous sections, but we use this different term for its well-established reference to veridical replay from an episodic storage, e.g., hippocampus, to a slow-learning system.

In addition, memory downscaling was implemented in the sensorimotor cortex, but not in the hippocampus, as the models were to update the hippocampal memory with new daytime memories in every cycle.

#### 2.2.1 Day-night performance oscillation in birdsong development

While experience replay is commonly used in continual learning [3, 24, 44], we found it the only mechanism that led to overnight performance growth (Fig 9A), which is incompatible with the real birdsong data. On the contrary, all the other models expressed overnight performance deterioration (Fig 9A). Therefore, in terms of being realistic, we can rule out experience replay as a candidate replay scheme. Nevertheless, we keep it in later analysis and discussion for its superb performance.

**Fig 9.**
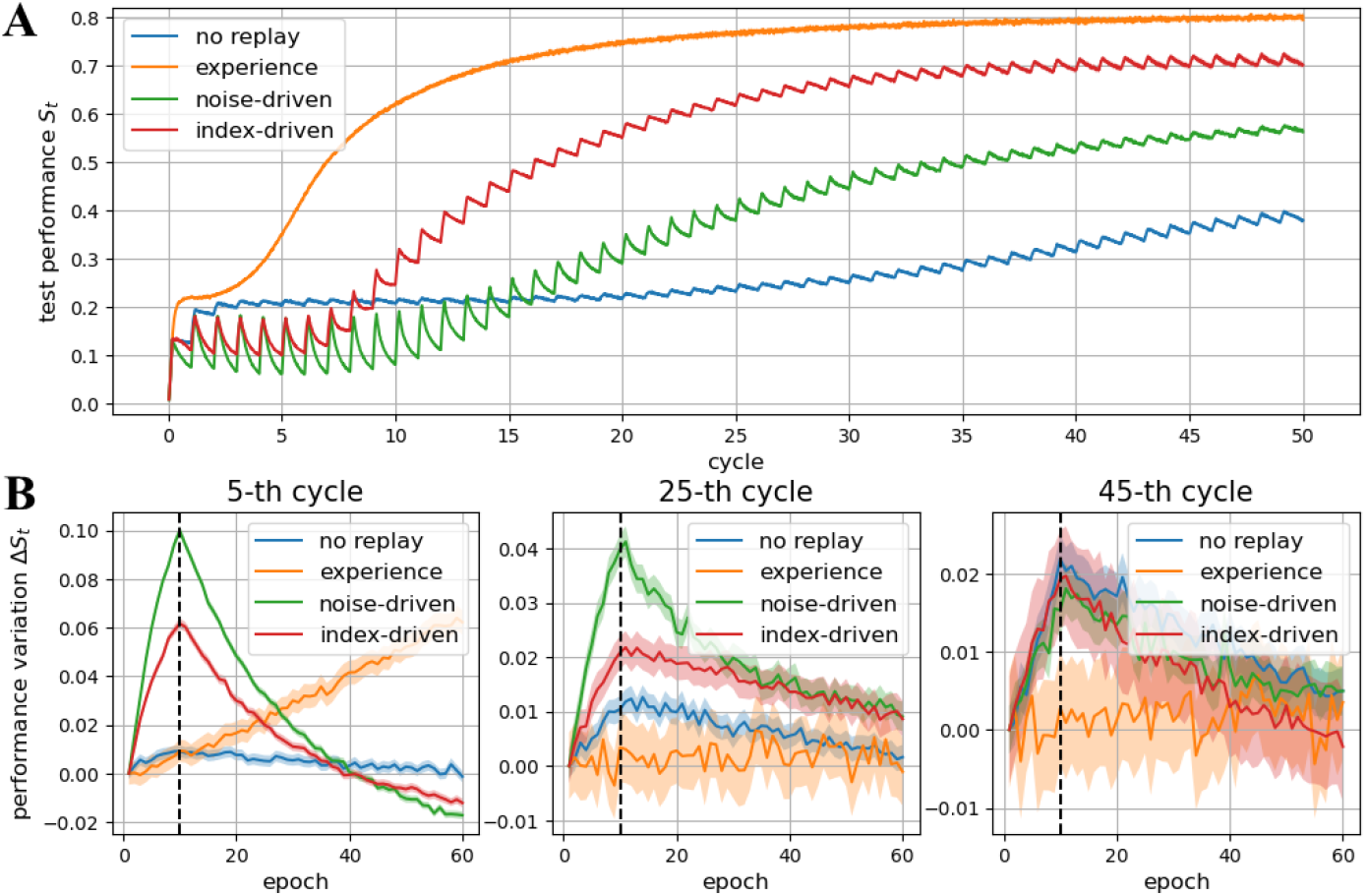
Performance growth and oscillation in development. While the 4 models’ overall performance increased in development throughout 50 day-night cycles (**A**), some of them manifested overnight performance deterioration, i.e., from the 10-th to the 60-th epoch in every cycle (**B**). Mean and 95% confidence interval from 100 trials are depicted.

While the overnight improvement of experience replay can be easily explained by its veridical replay samples, the overnight deterioration of other replay schemes, except no replay, cannot be attributed only to memory downscaling. With no replay, the performance oscillations kept increasing throughout the cycles, whereas, with noise- or index-driven replay, the learning trajectories appeared to oscillate more, at least in the early cycles (Fig 9B), but the overall performance grew rapidly beyond that seen in the no replay condition and was eventually closer to experience replay. Therefore, it were the replay samples that caused the different behaviours.

#### 2.2.2 Realistic developmental learning trajectories by replay

The questions of whether or not the performance oscillation observed in our models is realistic, and how this oscillation related to overall learning performance are intriguing. To answer these questions, we examined the magnitude of 3 types of performance changes across cycles (Fig 10A). The overall learning in development, i.e., the cycle-to-cycle performance gain, was most effective before, around, and after the 10-th cycle for experience, index-driven, and noise-driven replay, respectively, whereas it was ineffective and slow with no replay and downscaling only (Fig 10B).

**Fig 10.**
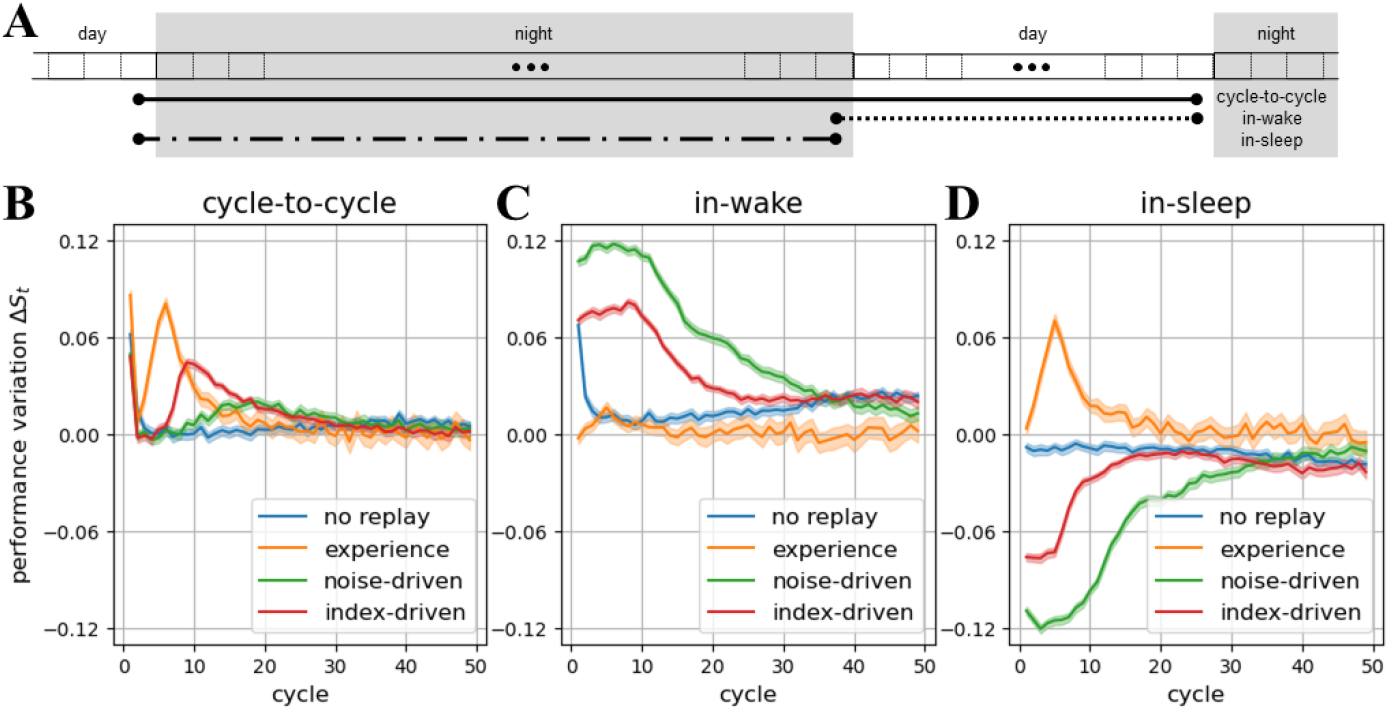
Performance change in day and night. The last epochs of day and night in every cycle (**A**) were used to calculated performance changes, from cycle to cycle (**B**), in a cycle’s wake phase (**C**), and in the sleep phase (**D**). Mean and 95% confidence interval from 100 trials are depicted.

Decomposing the cycle-to-cycle performance changes into in-wake and overnight changes, we observed the large gain in daytime and correspondingly the large deterioration in night in the noise- and index-driven models, and furthermore the magnitudes of these effects diminished across cycles (Fig 10CD). In contrast, although performance oscillation resulted in memory downscaling in the no replay model, its magnitude increased, rather than decreased, in the later cycles. Moreover, we note that the above observations were qualitatively consistent across different downscaling rates (Fig 19).

Following the idea in [16], we measured overnight performance deterioration and overall final performance, in order to determine if they were related. We assumed that individual zebra finches should share the same underlying replay mechanism but may differ in downscaling rates. For example, Fig 11A depicts the cumulative deterioration up to the 25-th cycle, and Fig 11AB depicts the average performance at the 25-th cycle, for different downscaling rates. It is clear from Fig 11C that the 2 measures were positively correlated in the noise- and index-driven models, but not in the others. As there was no natural choice of maturity epoch for our computational models, and this epoch may vary for the different models, we computed the correlation for all the epochs, and confirmed that the result held true for all epochs after the 20-th (Fig 11D).

**Fig 11.**
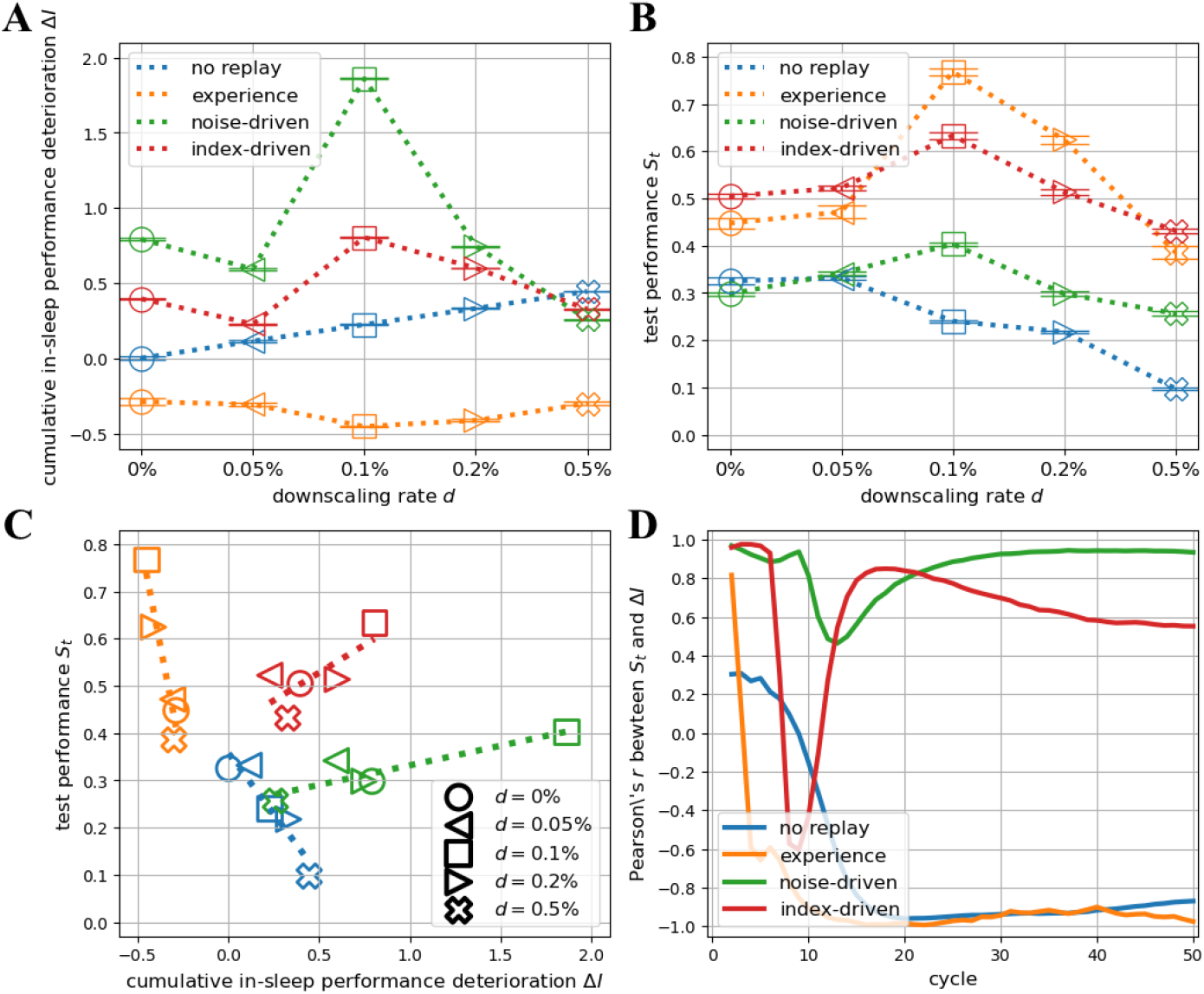
Relationship between final developmental performance and cumulative overnight performance deterioration. The effect of memory downscaling in the 4 models on the cumulative overnight performance deterioration, ∆*I* (**A**), and the final developmental performance, *S*_*t*_ (**B**), at the 25-th cycle. *S*_*t*_ is calculated by averaging the test performance of all the epochs in the cycle, and ∆*I* by adding up sign-flipped overnight performance changes up to the cycle. (**C**): The correlation between *S*_*t*_ and ∆*I* at the 25-th epoch for the different replay schemes and downscaling rates. The dahsed lines denote simple linear fits for different replay schemes. (**D**): The correlation between *S*_*t*_ and ∆*I* throughout development for different replay schemes. At every cycle, Pearson’s *r* is calculated from a simple linear fit (as in **C**).

Therefore, by comparing our simulated models to the real-life observations from birdsong development [16] (summarised by the bullet points in the beginning of the previous Section 2.2.1), we conclude that the noise- and index-driven models relying on sensorimotor cortical replay yield the most realistic oscillatory learning trajectories. The index-driven model is more favoured due to its better performance.

#### 2.2.3 Balanced and accurate replay benefits memory consolidation

Given our idealised modelling assumptions, replay (or no replay) is the only factor causing the behavioural differences in the models’ performance changes in development (Fig 10). With both no replay and experience replay, the magnitude of the overnight performance change could be simply explained by the current performance. The performance of the no replay model was reduced by memory downscaling, which was implemented as synaptic weight decay by a magnitude proportional to the weights (Section 4.1.4). Thus, the overnight deterioration became noticeable only after the performance was improved. On the contrary, experience replay provided the model with veridical samples that led to the immediate performance gain, and this gaining effect diminished as the overall performance approached the optimal plateau.

Similar explanations can also account for the other 2 models. Specifically, we can see from Fig 12 that the noise-driven model scored lower than the index-driven model in both evenness and exactness. These 2 metrics are extended but different from those used in Section 2.1.3, because we need to measure replay samples for all the syllables. While overall exactness is simply a weighted average of exactness of individual syllables, evenness is not some average of their frequencies. Instead, it evaluates how similar a replay distribution (of syllable frequencies) is compared to the uniform distribution. We note that evenness and exactness are conceptually akin to precision and recall (more details can be found in Section 4.2.3). For example, the replay samples generated by the noise-driven model were more biased towards certain syllables and worse at representing their true features, which explains the overall performance discrepancy between the 2 models.

**Fig 12.**
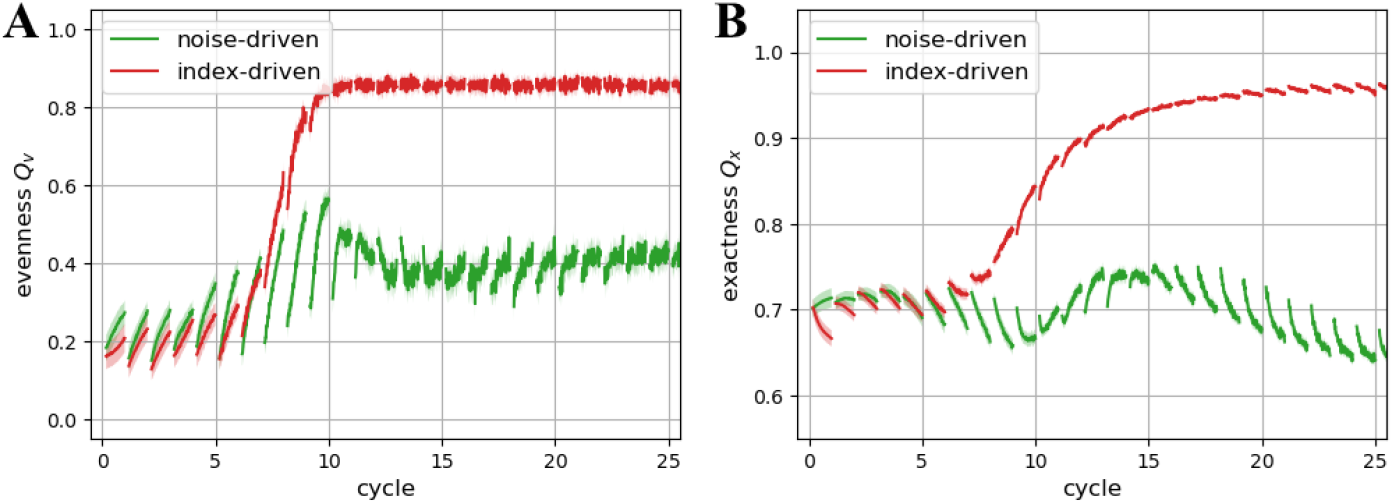
Quality of of the sensorimotor cortical replay samples in development. Evenness, *Q*_*v*_, measures how balanced replay samples were (**A**), and exactness, *Q*_x_, how accurate they were (**B**). The technical definitions of the two replay quality measures can be found in Section 4.2.3. Mean and 95% confidence interval from 100 trials are depicted. The experience replay model is not depicted, because its replay (from the hippocampus) did not vary across cycles.

Inspecting the changes in the replay measures and the performance on a nightly basis, we found replay evenness significantly better at explaining the overnight performance deterioration in the early nights, and exactness better at explaining this in the late nights (Fig 13). The changes were larger in the early cycles, and smaller in the late ones, because the sensorimotor cortical replay was gradually improved and stabilised in development. Consequently, the magnitudes of the deterioration in the noise- and index-driven models diminished across the cycles (Fig 10D).

**Fig 13.**
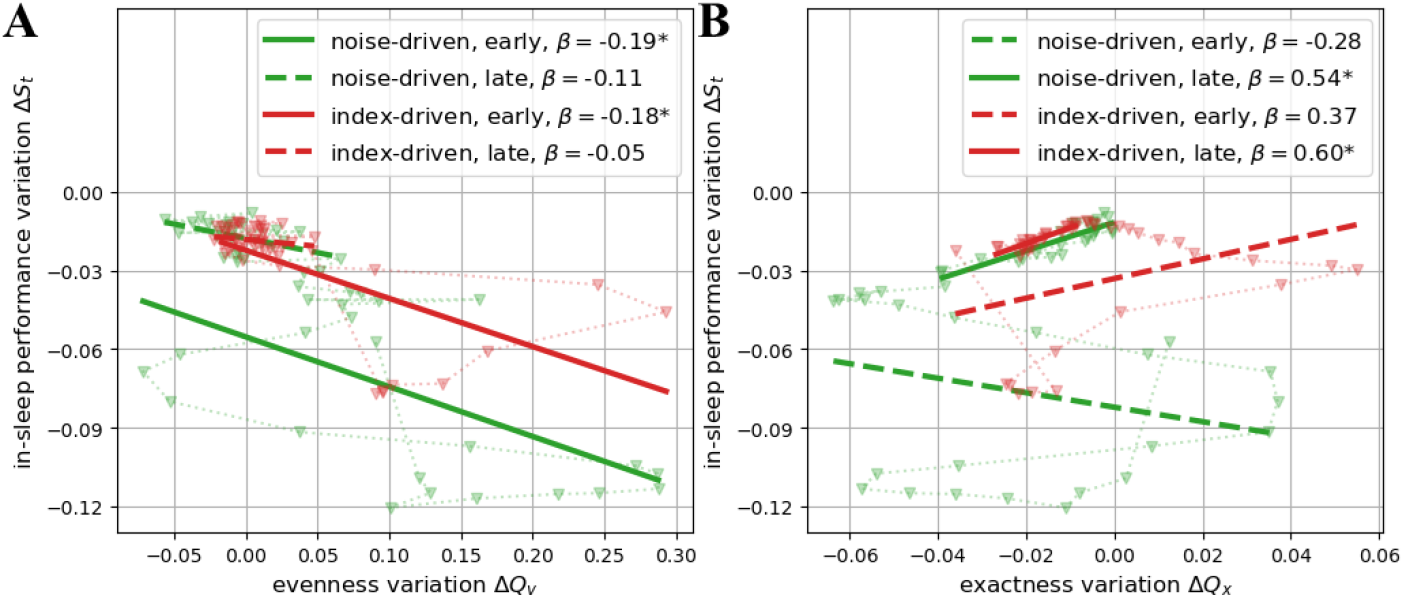
Correlation between the changes of performance and replay quality on a nightly basis. The triangular marker, connected in their temporal order, denote the same data as in Fig 12 for evenness (**A**) and exactness (**B**), respectively. The data before the 25-th night were grouped into the early epochs, and those after the late epochs. The thick lines of slope, *β*, were obtained by simple linear regression from the data. If the linear correlation is significant (*p <* 0.01), a solid line is depicted; otherwise, a dashed line depicted.

We note that, other than replay, the deterioration in the late cycles should probably be attributed more to downscaling. In addition, the early difference in replay exactness between the 2 models could not be well characterised by linear correlation (Fig 13B), but the early nights were the time when their replay quality measures and overall performance bifurcated (Fig 9A and 12).

Despite the overnight performance deterioration, the index-driven model achieved higher overall performance than the noise-driven model, due to its larger compensatory performance gain in subsequent wakeful learning (Fig 14A). The mechanism can be understood by considering consolidation via replay as effectively a pre-training process of a model before fine-tuning with true samples. We highlight the significant tradeoff between the 2 replay measures observed in the noise-driven model, but not in the index-driven model (Fig 14B). In other words, the noise-driven model was (statistically) bound to generate replay samples that were unfaithful to the truth, whereas there was no such constraint for the index-driven model. As a result, the index-driven model could consolidate noisy replay samples that were somewhat correlated to the true samples, benefitting the subsequent wakeful learning. We note in particularly that the increasing replay quality of the index-driven model in both measures (points in the first quadrant of Fig 14B) led to the relatively large day-night performance gain.

**Fig 14.**
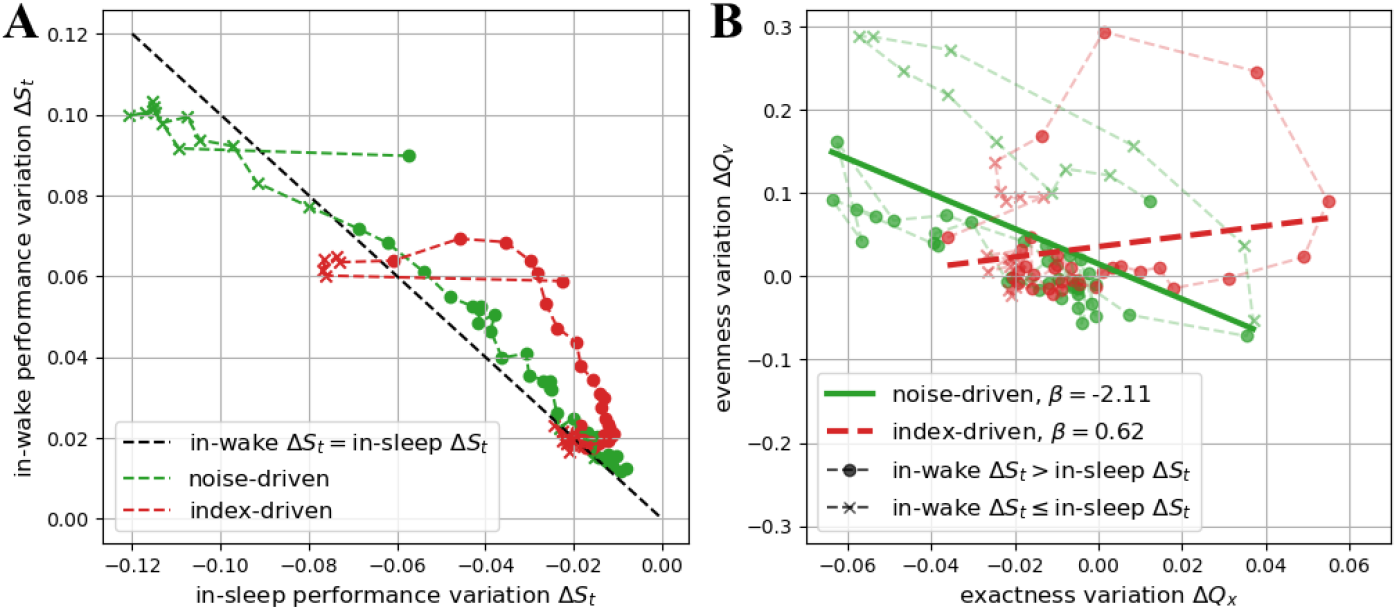
Replay quality modulates performance. (**A**): The relationship between the overnight and subsequent in-wake performance changes. Circles above the black dashed line denote larger performance gain in the wake phase than deterioration in the previous sleep phase, and crosses the opposite. (**B**): The changes in the two replay measures. The markers are chosen according to (**A**). The thick lines of slope, *β*, are obtained by simple linear regression from the data. If the linear correlation is significant (*p <* 0.01), a solid line is depicted; otherwise, a dashed line depicted.

## 3 Discussion

In this paper we investigated the roles of memory replay in two complementary learning systems (CLS) models capturing realistic sleep effects observed in real-life targeted memory reactivation (TMR) in Sections 2.1.1 to 2.1.3, and birdsong development experiments in Sections 2.2.1 to 2.2.3. The two CLS models are identical in architecture, consisting of an idealised hippocampus modelled by a stochastic simple memory (SSM) and an idealised neocortex (or sensorimotor cortex) modelled by a restricted Boltzmann machine (RBM).

Other than the differences in the learning and the replay protocols specified by the two simulated experiments, the two models are distinct in terms of where replay is generated. While the TMR model produces replay samples using its hippocampus, the birdsong model does so using the sensorimotor cortex.

### 3.1 Insights from targeted memory reactivation (TMR)

We found that memory consolidation in the neocortex via hippocampal replay could account for the realistic, primary TMR effects, i.e., the targeted memory was strengthened for a prolonged time. In addition, the memory gain was not obtained at the cost of other memories, unless the hippocampus was highly susceptible to external noise. This secondary effect implies a criterion for filtering subjects, in order to minimise the negative effects of TMR manipulations. It might also explain how traumatic memories can be mitigated by the TMR technique.

Although the replay biased by the cues was ultimately responsible for the post-cueing TMR effect, we did not observe significant post-cueing changes in the replay measures per se, except in the plastic and noise-driven model which yielded significant performance improvement (Fig 7). We thus consider that the slow, statistical learning feature of the RBM accounts for the effect, as this feature probably made the model performance more sensitive to the samples and their biases encountered early in learning, particularly during the cueing periods. Even with balanced training samples as in the birdsong model, the RBM took many epochs before its memory became unbiased (Fig 12A).

We note that it is difficult to map the number of sleep epochs in our simulations linearly to sleep time in real-life experiments, especially given the fact that multiple distinct sleep stages exist, and they are believed to be responsible for different brain functions [26, 45, 46]. Our simplified assumption that a model constantly transfers knowledge from hippocampus to neocortex is more suitable for modelling slow wave sleep [7, 8, 26, 35], whereas other computational models showcase the possibility that the neocortex can perform more advanced memory processes, e.g., incorporating replay in other sleep stages, so that memory consolidation is more effective and robust [47, 48]. Moreover, whether the offline processing epochs are interpreted as ‘sleep’ is entirely up to a modeller, because replay also occurs in wakefulness [33, 49]. The model can be extended to address these concerns, e.g., by implementing sleep cycles, potentially interleaving different replay mechanisms, or day-night cycles, in which a model repeats learning and replay as in our birdsong model.

### 3.2 Insights from birdsong development

We show that, amongst our 4 types of replay, the sensorimotor cortical replay driven by hippocampal sequential signals (i.e., index-driven replay) produces the most closest fit to the characteristics of learning observed across development of songs in juvenile zebra finches who replay these songs each night, i.e., the magnitude of the performance oscillation decreased gradually across multiple nights, and the overall overnight performance deterioration correlated positively to the final developmental performance.

We highlight that such index-driven replay, rather than the noise-driven replay, could lead the model to reach comparable performance to experience replay (i.e., veridical hippocampal replay) in the late development. As suggested by [16], lack of true auditory feedback in sleep could explain overnight performance deterioration, but it might also make replay more exploratory, contributing to better performance in the long term. Similarly, there was no true feedback for our models with either the noise- or the index-driven replay. It was important, however, to somewhat control the stochastic process of generating replay samples, e.g., hippocampal signals in the index-driven replay, which still encouraged the RBM to adjust its weight configuration away from spurious, local optima, but not too much from the global optimum.

In contrast, experience replay was always the first to demonstrate a rapid performance growth in the early development, despite being overtaken by the index-replay under some conditions (Fig 19). Since the index-driven replay allows a model to reconstruct samples of high quality while saving the resources of storing veridical data (given the sensorimotor cortex has been properly trained), we suggests that it can actually be more beneficial (in addition to being natural) for a hippocampus to store and replay new memories as veridical episodes, but to memorise old memories in a fragmented manner and to trigger neocortical replay using these fragmented signals. As these signals become more abstracted and generalised due to learning and forgetting, they might perform the role of index as idealised in the hippocampal indexing theory [41, 43].

### 3.3 Different roles of experience and generative replay

While we named the various replay schemes to accommodate the different contexts of the two real-life experiments, we reiterate that experience replay in the birdsong model is the same as spontaneous replay in the TMR model, as in both cases veridical episodic replay samples were produced by the hippocampus idealised as an SSM. In addition, we note that implementing noise-driven hippocampal replay would make tiny differences compared to experience replay in the birdsong model, because the hippocampus keeps updating its memory with new training samples in the daytime epochs. However, such tiny differences could accumulate and amplify, resulting in the performance discrepancies observed in the TMR model.

Although it has been considered biologically less plausible, experience replay is commonly used in continual learning to mitigate the problem of catastrophic forgetting [3, 24, 44]. An alternative, generative replay, has caught more attention of the ANN community only recently due to increasing concerns over computational and ethical restrictions [50]. The sensorimotor cortical replay which we implemented in our birdsong model belongs to this type, as the RBM (or more generally any generative model) was able to maintain traces of old memories after proper training [40, 51–53].

Since the RBM performed slow, statistical learning, the quality of its replay samples would not be high until it was properly trained, leading us to make the suggestion in the previous Section 3.2. More importantly, the birdsong model showed us the difference that generative replay could make even before the RBM was well trained. Thus, we cannot rule out the possibility of some unknown coordination between the hippocampus and the medial entorhinal cortex, even though their replay patterns were reportedly uncorrelated [10].

We consider all our implementations of various replay mechanisms representative and idealised with respect to different hypotheses within the theoretical framework of system consolidation [1, 18, 54], but in reality they are unlikely to be exclusive to one another. Comprehensive reviews on the aforementioned and more replay schemes studied in ANNs can be found in [55, 56], and reviews on their inspirations, connections and implications regarding biological systems can be found in [1–3].

### 3.4 Differential effects of downscaling and replay

Memory downscaling is widely used in ANNs, and can be roughly divided into weight decay [57, 58] and pruning [59, 60]. Although we consider them to be plausible implementations subscribing to the Synaptic Homeostasis hypothesis [61, 62] as in [63, 64], they are largely simplified from the biological reality, and other implementations also exist [65]. Notably it is often unnecessary for ANNs to undergo sleep periods, and downscaling such as weight decay is typically integrated into their training protocols [66], which is believed to bring in the same benefits as if performing downscaling in a dedicated sleep period [63].

We highlight that downscaling can perform a role which is complementary role to those of replay and consolidation. Distinctively in our TMR model, the plastic and noise-driven hippocampus produced better replay samples in general, not only for the targeted memory (Fig 18). This model thus outperformed the others in the long term regardless of the conditions (Fig 3).

Interestingly, it would be impossible for the hippocampus to maintain the episodic memories with constant replay and consolidation but no downscaling. Memory downscaling effectively time-stamped the memories, as the older ones (including those originally learned and those later consolidated) must have undertaken longer retention period and tended to downscale more. It was thus more likely for the SSM to remove memories evenly across all categories (given balanced training samples). On the contrary, without downscaling, it would be equally likely for the model to obliterate the same number of memories from a single category as from multiple categories, downgrading the replay quality faster.

Regarding the relative failure to maintain the episodic memories in the other models, we note that the episodic memories were eroded in the two stable models due to downscaling only (but no consolidation via replay), and that the replay samples were increasingly biased in the plastic and spontaneous model. To summarise, replay and downscaling can cause memory loss on their own, but jointly they slow down forgetting.

We also note in the birdsong model the effects of downscaling on performance deterioration and final performance were nonlinear given replay and consolidation (Fig 11AB). How biological or ANNs optimise for the tradeoff between replay and downscaling is however beyond the scope of this paper.

## 4 Methods

### 4.1 Computational models of learning systems

While emulating learning systems, e.g., real brains, could be pursued by focussing on building models of their structures, we approached the same goal from an alternative perspective concerning primarily the universal functions of any (useful) learning systems, that is, memory acquisition (learning), retention (storage) and retrieval (recall). This approach permitted the nearly identical architectures used for our models of human and bird brains, despite any anatomical differences of the real brains. More importantly and conveniently, we could define memory replay for any learning system to be the process of memory recall followed by consolidation (learning) of recalled memories.

Explicitly, we consider only artificial neural networks (ANNs) for our models, and we can write down an ANN’s recall function as

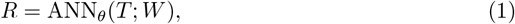

where *T* denotes an input presented to the ANN, *R* its output, i.e., the response, and *θ* its intrinsic parameters. We highlight that the memory of this (or any arbitrary) ANN is assumed to be encoded fully in its weights, *W*, and the changes, ∆*W*, given *T*, can be obtained by the ANN’s learning function,

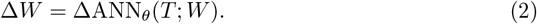

While normally *T* represents training or testing data and *R* the responses, in the context of memory replay *T* should be interpreted as input signals triggering replay and *R* the replay samples. Consequently, the ANN performing memory consolidation using its own replayed samples updates its weights by

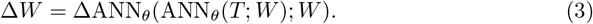

Moreover, if multiple (sub-)ANNs communicating with one another (in a whole-brain model) are considered, Eq (3) can be modified and extended according to their interactions. For example, in a model composed of two ANNs, replay from model 1 may be consolidated by model 2, that is,

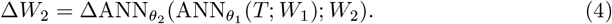

As in many ANN works, we use *replay* to mean reactivation of a set of memory patterns in our models, not necessarily following a temporal order. It is considered almost interchangeable with *reactivation* [21] and *rehearsal* [44] from an abstract perspective of computational modelling, even though these terms can be associated with different phenomena in the neuroscience literature.

#### 4.1.1 Complementary learning systems (CLS)

Complementary learning systems (CLS) is a theoretical model proposed by [21] primarily for explaining how a mammalian brain integrates different learning and memory systems to acquire, represent, and apply knowledge effectively. In its most classical form, the CLS theory asserts that the brain has a fast system and a slow system performing complementary roles in learning and memory. Specifically, the fast system is the hippocampus, capable of rapid learning of specific episodic experiences or events. It creates sparse, pattern-separated representations, allowing for the encoding of detailed memories. The slow system is the neocortex, responsible for slowly integrating and compressing the knowledge acquired from the hippocampus into more structured, overlapping representations. It extracts statistical regularities and forms generalised knowledge. In particular, the hippocampus uses replay to transfer knowledge to the neocortex.

Over the decades, the CLS theory has been well studied and modernised via empirical evidences [24, 25], and has become a modelling framework that inspires many artificial learning and memory systems, especially ANNs, to implement replay-like algorithms [1] to mitigate catastrophic forgetting [22] (or the stability-plasticity dilemma [23]), which refers to the high risk of old memories being eroded or even erased by new memories in ANNs.

While an avian brain does not have hippocampus or neocortex as in a human brain (in a TMR experiment), there exist functionally similar brain structures, e.g., hippocampal formation, hyperpallium [11–14]. Considering additionally the critical role of replay in zebra finches’ song development [15], we are inspired to emulate both human (mammalian) and avian brains by a CLS model consisting of two ANNs. Specifically, the fast system is considered to be embodied by a *stochastic simple memory* (SSM) [28, 29], and the slow system by a *restricted Boltzmann machine* (RBM) [30, 31] (Fig 1).

We consider the SSM an idealised model of the CLS hippocampus (as in [17, 67, 68]), because it can store and recall data in a rapid, veridical, and pattern-separated manner as if episodic memories. Notably the model is a typical exemplar model, in addition to being a connectionist model. It is almost identical to the REMERGE model [69], except that we do not consider recurrent activation between the two layers.

We consider the RBM an idealised model of the CLS neocortex (as in [17]), because its weight matrix *W* statistically extracts information from a large number of training samples, and encodes memories implicitly as prototypes.

#### 4.1.2 Restricted Boltzmann machine (RBM)

The restricted Boltzmann machine (RBM), initially proposed under the name Harmonium by [30], is an ANN with a characteristic architecture consisting of two sets of neurons (see Fig 1). The feature (visible) neurons with activities, *v*_*i*_ ∈ *{*0, 1}, *i* ∈ *ℳ* = {1, 2, …, *m*}, can observe external data, but the latent (hidden) neurons with *h*_*j*_ ∈ {0, 1}, *j* ∈ *𝒩* = {1, 2, …, *n*}, can only communicate with the visible ones only via synapses, *w*_*ij*_ ∈ ℝ. All the synapses are assumed to be bidirectional and symmetric in terms of their weights, i.e., *w*_*ji*_ = *w*_*ij*_, forming a weight matrix *W* = [*w*_*ij*_]_(*i,j*)∈ *ℳ ×𝒩*_.

As there is no lateral communication within each set, *h*_*j*_ is completely (but stochastically) determined by *v*_*i*_. More specifically,

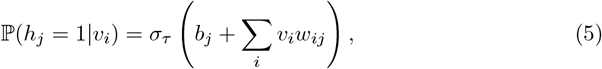

where

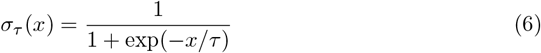

is the sigmoid activation function with *τ* modulating the randomness level (fixed to be 1 in this work), and *b*_*j*_ denotes the intrinsic bias of *h*_*j*_. Following a nearly identical formula to Eq (5), *v*_*i*_ is completely determined by *h*_*j*_ via ℙ (*v*_*i*_ = 1|*h*_*j*_). Consequently, given an input vector *T*, the RBM’s response is

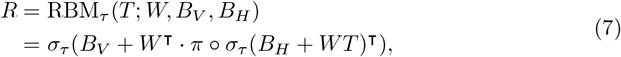

where *B*_*V*_ = [*b*_*i*_]_*i*∈ℳ_, *B*_*H*_ = [*b*_*j*_]_*j*∈𝒩_, ^⊺^ and *°* denote matrix transpose and function composition, respectively, and *π* randomly binarises the probabilities obtained via *σ*_*τ*_.

The weight matrix *W* is statistically learnable using the contrastive divergence algorithm [31]. Specifically, we use the algorithm with one iteration of Gibbs sampling with a fixed learning rate, *γ*, which gives

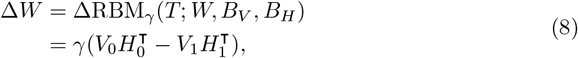

where *V*_0_ = *T, V*_1_ = *R* and *H*_*k*_ = *π ° σ*_*τ*_ (*B*_H_ + *WV*_*k*_)^⊺^. We note that the biases, *B*_V_ and *B*_H_, are updated by the same rule, as they are mathematically equivalent to *W*.

Due to the statistical nature of the RBM, a small learning rate is required, and we set *γ* = 0.01. Readers are referred to [66] for more details on the RBM and its learning algorithm.

#### 4.1.3 Stochastic simple memory (SSM)

The stochastic simple memory (SSM) is our ad hoc implementation of Marr’s archicortex model, namely *simple memory* [67, 68]. In particular, we view it as an ANN that has an architecture similar to the RBM, consisting of feature and latent (codon) neurons. It has two distinct features. Firstly, the activities of its hidden neurons influence one another via lateral inhibition (see Fig 1), promoting an extremely sparse (nearly one-hot) activation pattern. Secondly, the SSM learns by replacing the weakest memory with the new one.

Specifically, when the SSM with a weight matrix, *W*, receives an input vector *T*, the latent activity is

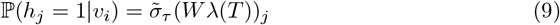

where

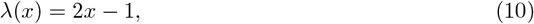

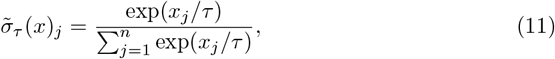

and consequently the response is

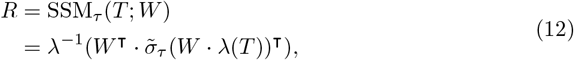

where *λ*^−1^ is the inverse transform of *λ*. We note that the linear transform *λ* merely serves the function of unifying two conventions of encoding presence and absence of a feature, {1, 0} vs {1, −1}, and 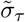 is the softargmax function, with *τ* tuning the randomness level. If *τ* = 0, the softargmax function reduces to the argmax function, and consequently only the hidden neuron whose weights are the most similar to the input (maximising ∑ _*i*_ *λ*(*v*_*i*_)*w*_*ij*_) is activated. If 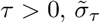 yields a vector of probabilities adding up to 1. Stochastic modelling of stimulus-response choice by the softargmax function (11) appeared no later than [28], with similar ideas proposed repeatedly [70, 71], and it has become a common practice for contemporary ANNs to use the function for categorisation.

In learning, the weight matrix *W* is updated by changing all the synaptic weights connected to the hidden neuron of the weakest activation to the pattern of an input vector *T*, that is,

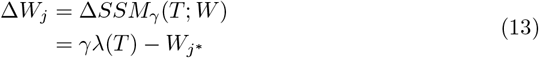

where 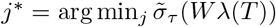. This algorithm guarantees 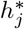 to become the maximum (given sufficiently large *γ*) if *T* is re-presented after learning *T* until being forgotten.

As learning (and forgetting) an item affects a single memory (encoded by a single hidden neuron) at a time, the learned memories are episodic and pattern separated. As such, we set *γ* = 1 to maximise the SSM’s learning efficiency. While the response from Eq (12) is a weighted average of all memories, a small *τ* (fixed to be 1 as for the RBM) ensures the response to closely resemble one of the episodic memories.

#### 4.1.4 Replay schemes

As stated earlier, an ANN is assumed to produce replay by its recall function (1). Consequently, the SSM and the RBM do so by Eqs (12) and (7), respectively. Depending on the nature of replay and the replay cue (i.e. the input *T*), we consider four replay schemes for our models:

- *No replay*: There is no *T* and no replay. This scheme offers a baseline for comparisons.
- *Cue-* or *index-driven replay*: *T* encodes (partial) information of training data. Replay samples are expected to resemble the corresponding training samples.
- *Noise-driven replay*: *T* is completely random. Replay samples are expected to be highly dependent on a model’s memory.
- *Spontaneous* or *experience replay* (for the SSM only): *T* does not exist or equivalently *λ*(*T*) = 0. As all the hidden neurons are (in)activated at exactly the same level, an episodic memory is randomly chosen.

We dismissed spontaneous replay for the RBM, as it yielded ill-behaved replay samples according to our early-stage experiments.

#### 4.1.5 Downscaling schemes

Memory downscaling, or generally weakening synapses in biological nervous systems, can be beneficial, because it contributes to the restoration of synaptic homeostasis, reducing metabolic costs and signal saturation [61–63, 65]. It also improves generalisation of ANNs [72], including RBMs [66], by moderating overfitting, and has become widely used with various implementations in machine learning [57, 58].

In total we assumed three downscaling schemes:

- *Synaptic decay* scales down the weight matrix *W* by a decay rate *d*_sd_ ≥ 0,

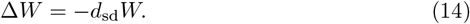
- *Synaptic pruning* removes non-zero elements smallest in magnitude in *W* by a percentage of *d*_sp_, and replaces them by zeros.
- *Neuron pruning* removes memory traces in *M* or hidden neurons in *W* with the smallest L1-norms by a percentage of *d*_np_, and re-initialises them with random values.

For simplicity, we fixed all these rates be equal to the downscaling rate *d*, i.e. *d*_sd_ = *d*_sp_ = *d*_np_ = *d*, implemented these schemes in all our models.

As we set *d* ≤ 1% in all the simulations,^2^, only the synaptic decay rule could change model performance significantly. It was too small for (synaptic and node) pruning to cause noticeable effects [59, 60]. All the model stay unchanged after downscaling if *d* = 0.

Due to downscaling, older memories that is usually weaker in the SSM are more likely to be obliterated when it acquires new memories.

### 4.2 Simulated experiments and data analysis

#### 4.2.1 Synthetic data in experiments

In this paper we conduct numerical experiments with synthetic data. Any training or test sample *T* are instantiated from a category prototype *C* (e.g., an item pair or a song syllable) that are randomly generated before every simulation trial. Formally, we make the following assumptions:

1. 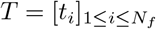 and 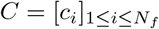 are vectors representing *N*_*f*_ measurable features;
2. For any *i, t*_*i*_ ∈ {0, 1}, where 1 means the presence of the *i*-th feature, 0 the absence of it;
3. For any *i, c*_*i*_ ∈ [0, 1], which denotes the probability of the *i*-th feature’s presence;
4. Every category is fully and uniquely characterised by its category prototype *C*;
5. Any sample *T* associated with category *C* is instantiated by a sampling process *π* which randomly binarises all *N*_*f*_ features of *C*.

Specifically, for any *i*,

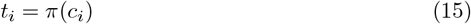

performs a Bernoulli trial, and

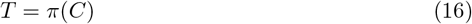

is composed of *N*_*f*_ independent Bernoulli trials.

We randomly generate *N*_*c*_ category prototypes at the start of each simulation trial. While our models can encounter training samples from the *N*_*c*_ categories in an uneven manner, they are tested by samples generated uniformly from all the categories.

Notably, every *T* is uniquely associated with one and only one *C*, but true category labels are always inaccessible to our models, as they perform unsupervised learning. Although it is possible that *T*_1_ produced by *C*_1_ and *T*_2_ by a different category *C*_2_ are similar to each other, the probability of of such events diminishes geometrically with respect to the number of features (at the order of 10^2^ in this work), and is thus assumed negligible in all our analysis.

Due to our focus on replay by cueing (or indexing), which is inspired by cued recall, we consider any feature vector composed of two regions, a recall region and a cue region. Specifically for a category prototype, we define

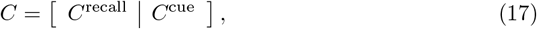

which concatenates features in the two regions. A cueing sample from a category is instantiated from a modified prototype,

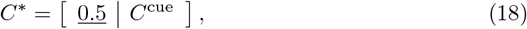

where <u>0.5</u> denotes a vector whose individual elements are all 0.5. Test samples are obtained in the same manner.

In Sections 2.2.1 to 2.2.3, a bird song is conceptualised as a sequence of a handful of syllables, which are represented by feature vectors of prototypes. It can also be written as the matrix (17), where *C*^recall^ and *C*^cue^ = *C*^index^ are assumed to encode the acoustic and the temporal information of the song. Moreover, we assume that temporal context vectors encode orthogonality (i.e., pseudo-sequentiality) in the following explicit form,

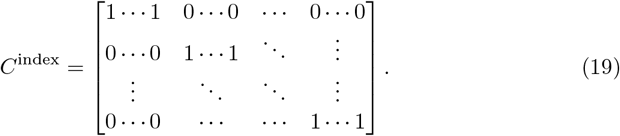

In Sections 2.1.1 to 2.1.3, the recall region is further divided into a visual-cue region for probing and a visual-recall for measuring test performance, as TMR cueing is an experimental manipulation that is no more presented in test, ensuring any performance discrepancy between conditions is indeed a memory effect. The matrix (17) becomes

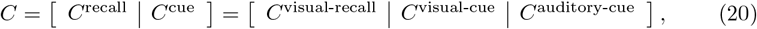

where the auditory-cue region encodes TMR cues, and

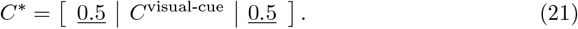

#### 4.2.2 Metrics for test performance

In this paper we choose cosine similarity as the primary metric measuring similarity between vectors. We do not report results based on other similarity metrics, e.g., L1 norm, Rand index, because we do not observe qualitative differences. Which metrics are suitable for a specific research field [73, 74] and how to maximise our models’ quantitative performance are beyond the scope of this paper.

Specifically, we always deploy a cued recall test, in which we probe our models with a test sample *T*, instantiated from *C*^*^, whose recall region is completely randomised as in Eq (18) or (21) (Section 4.2.1). Responses from the models *R* are then compared to true prototypes by

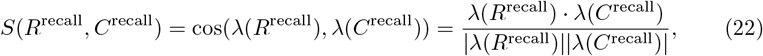

where *λ* is a linear mapping acting on all features,

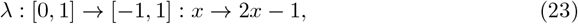

and | · | denotes the Euclidean length of a vector. *λ* effectively shifts the centre of feature vectors (whose elements ranging between 0 and 1) to 0.5 and doubles their length (which has no effect on the cosine similarity), as we assume that 1’s and 0’s in a feature vector are symmetric to each other. Notably this assumption can be unrealistic, especially when *x* is considered to be a pre-processed neural signal which is commonly sparse coded in real brains.

The average similarity between a category prototype *C* and all 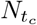 responses *R* obtained from a model when it is probed by test samples generated from the category is considered the metric for category test performance,

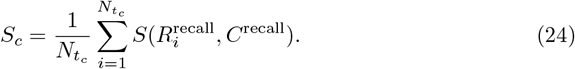

Similarly, the overall test performance is calculated by averaging similarity values over all such (*R, C*) pairs, that is,

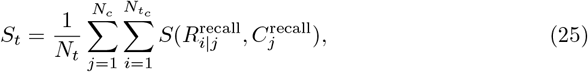

where 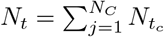 is the total number of test samples.

The chance level of test performance *S*_*t*_ and *S*_*c*_ is nearly 0. Firstly, it can be mathematically proven that cosine similarity of two independent random vectors diminishes uniformly to 0 if their dimension increases to infinity. Next, a model without useful memory (and thus yielding chance-level performance) would produce random *R*^recall^ independent of *C*^recall^. As we run simulations with reasonably large dimension (*N*_*f*_ = 200 or 300), the chance-level performance is indeed negligible.

When measuring the performance of a CLS model, we consider only *R* from the RBM.

#### 4.2.3 Metrics for replay quality

In this paper our models are designed to produce replay samples (Section 4.1.4), which are denoted by *R*. It is impossible in the schemes of experience replay and noise-driven replay to measure their quality by the similarity metrics *S*_*t*_ or *S*_*c*_ in Eq (24) and (25), for they relying on pairwise comparisons between *R* and *C*, but *C* not provided by these two schemes.^3^ We thus propose two replay quality metrics, exactness *Q*_*x*_ and evenness *Q*_*v*_, which are defined to be

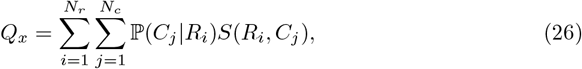

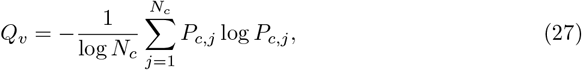

where ℙ (·) denotes a probability, ℙ (·|·) a conditional probability in particular, and log(·) the logarithmic function, and

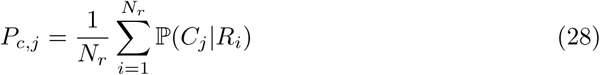

describes the marginal distribution of category *j*.

These definitions follows the idea of precision and recall, which has been proposed to measure the sample quality produced by generative models [50, 75, 76]. An ideal memory model would be capable of reconstructing samples of high fidelity from all categories. In practice memories may interfere with one another, and thus there is a tradeoff between generating accurate samples (high exactness) and recalling all categories (high evenness).

The specific implementation is different from those in [50, 75, 76], because we are not considering realistic images. We choose *Q*_*x*_ to be based on cosine similarity for consistency with the test performance metrics. However, every replay *R*_*i*_ can be associated with multiple categories of different weights ℙ (*C*_*j*_|*R*_*i*_). We define evenness *Q*_*v*_ to be Pielou’s evenness, initially designed for measuring biodiversity [77], as it ranges between 0 and 1 for 0 implying that all replay samples belong to a single category and 1 a uniform replay distribution. A model is expected to yield a decent test performance across all categories when both its replay quality indices, *Q*_*x*_ and *Q*_*v*_, are close to 1.

In addition, as we know the ground truth values of our synthetic data, we can calculate the metrics exactly rather than conducting estimations. Specifically, ℙ(*C*_*j*_|*R*_*i*_) can be calculated by the Bayes theorem, and *S*(*R*_*i*_, *C*_*j*_) by Eq (22).

#### 4.2.4 Data generation and analysis

Only synthetic data was used. All statistics were computed using the standard numpy and scipy packages in Python. The solid curves in Fig 4-7 were smoothed from the data points by a Savitzky–Golay filter of order 3 and sliding window width 11.

## Acknowledgments

This work was supported by the ERC grant SolutionSleep, 681607, to PL.

## Extended figures

**Fig 15.**
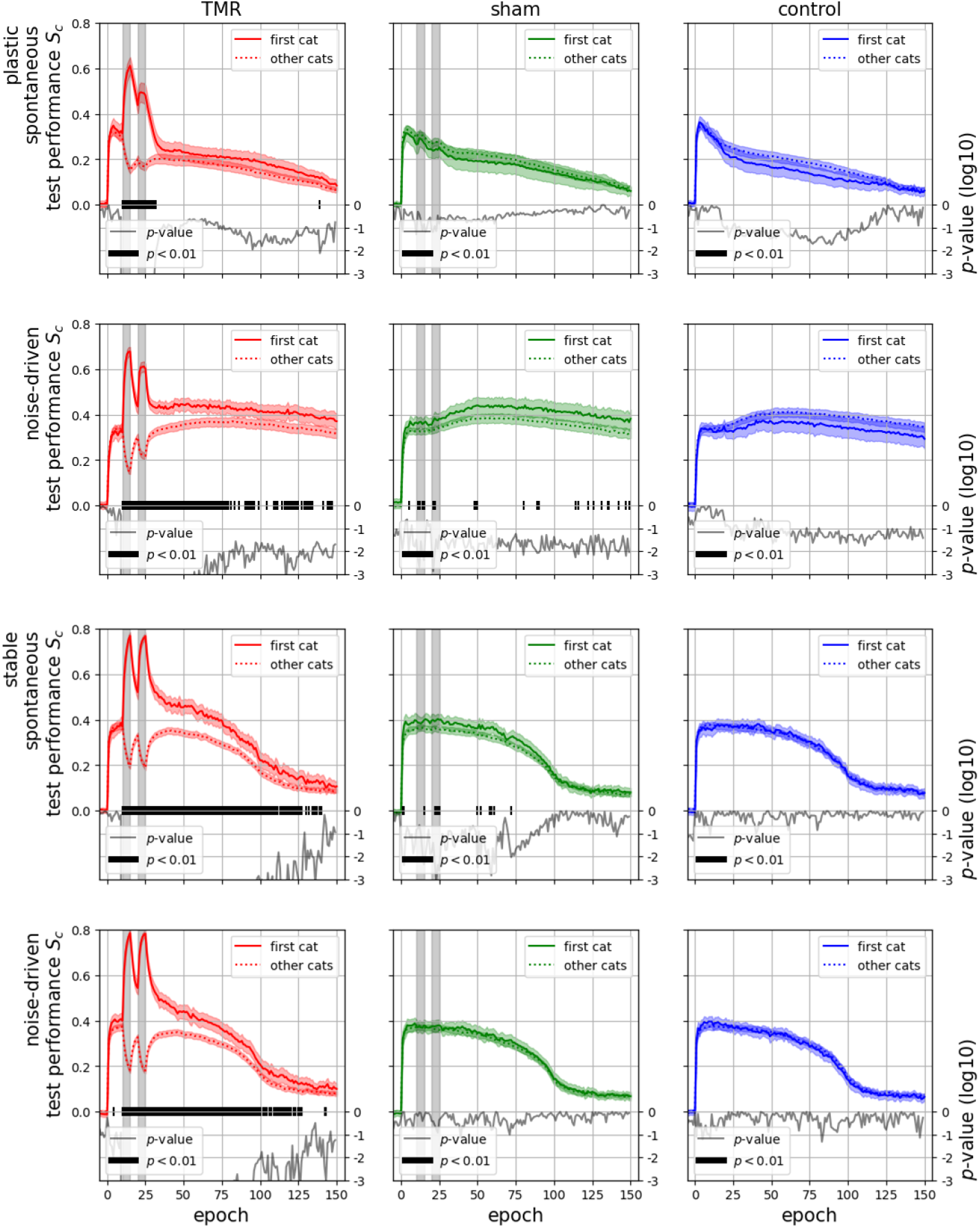
Only the TMR manipulation introduced performance discrepancies betweeen the first category and the others. Under the sham and the control conditions, there were no significant differences between the first category and the others. Therefore, their performance was merged into overall averages as in Fig 3.

**Fig 16.**
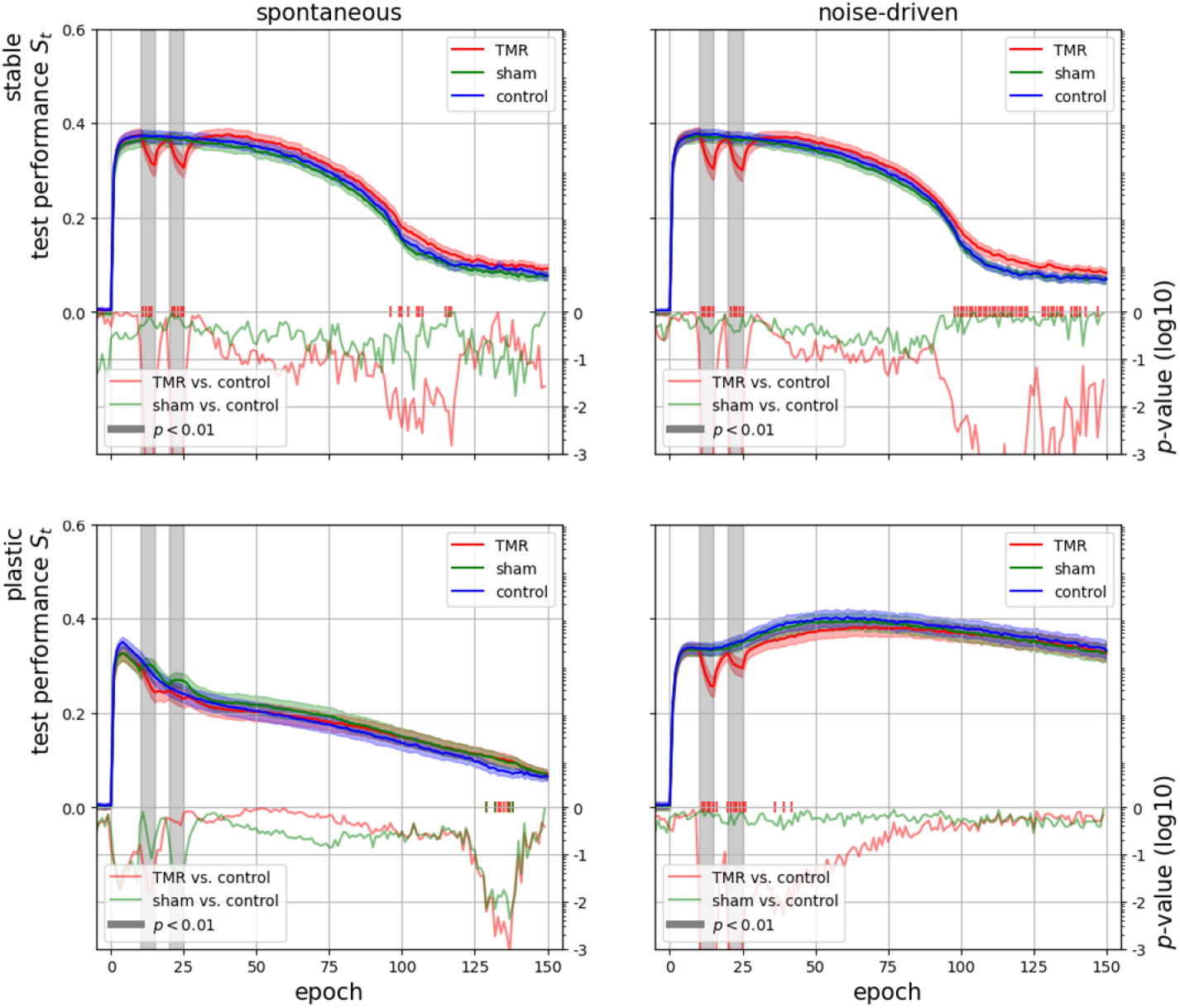
Only in some post-cueing brief time periods, the TMR manipulation resulted in significant discrepancies in the overall performance. This was probably due to the fact that in total 5 categories were to be learned, and significant performance discrepancies in the targeted memory was not large enough to affect the overall performance.

**Fig 17.**
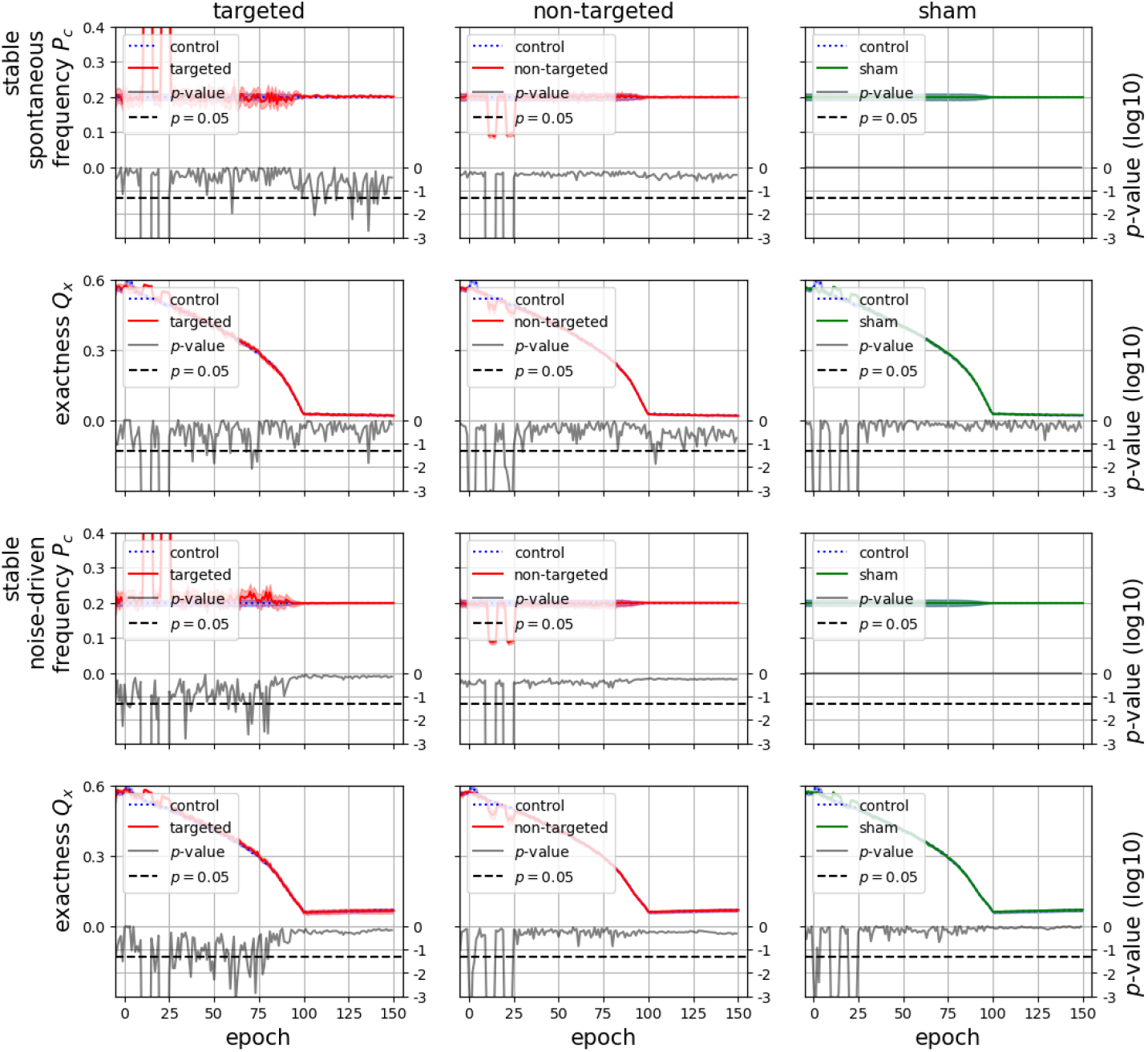
Complete replay measures in the two stable models. Only in some brief post-cueing time periods under the TMR condition, the replay frequency *P*_*c*_ and the exactness *Q*_*x*_ of the targeted memory were significantly different from the control (*p <* 0.05).

**Fig 18.**
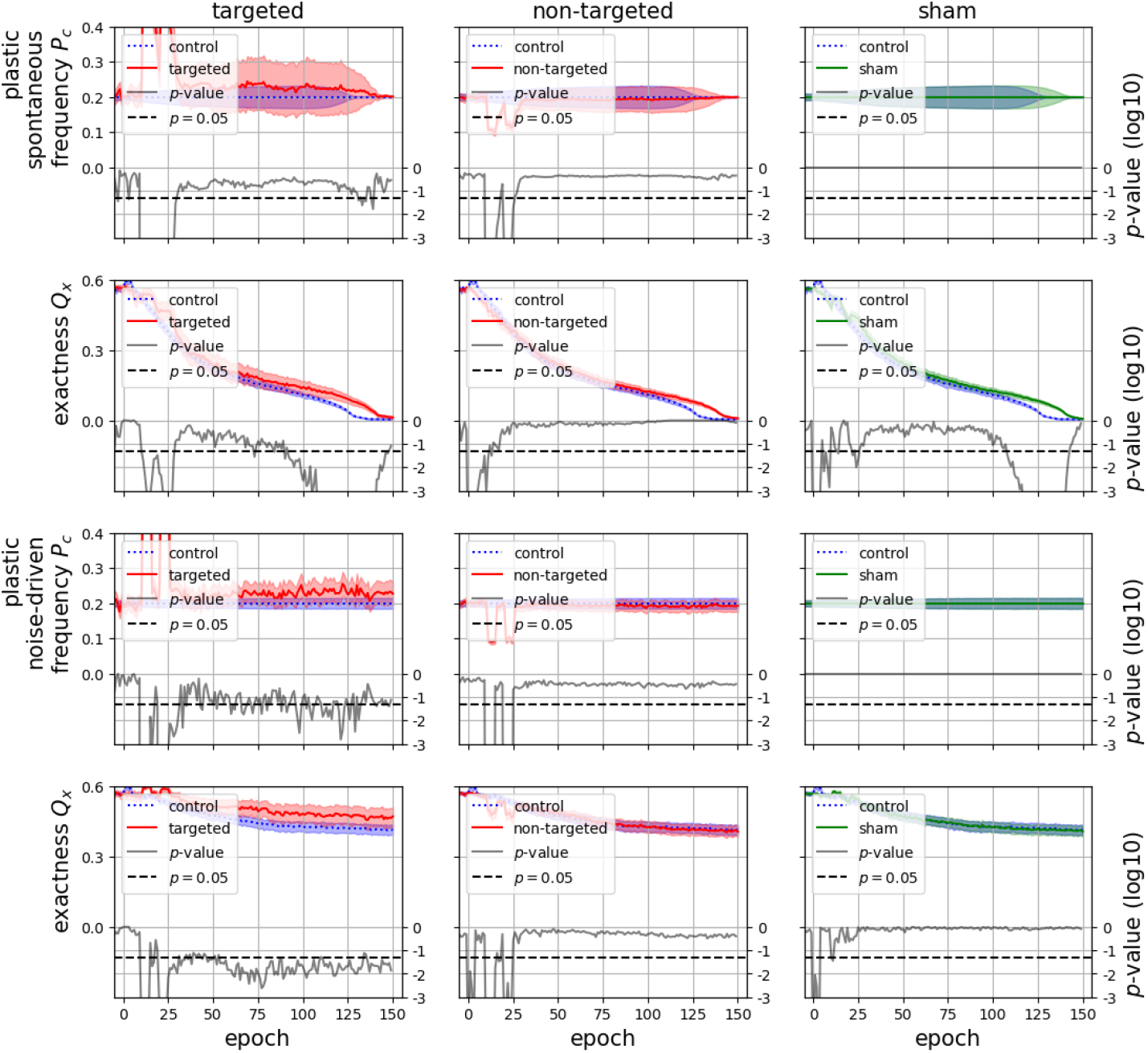
Complete replay measures in the two plastic models. Under the TMR condition, the replay frequency *P*_c_ and the exactness *Q*_*x*_ of the targeted memory in the plastic and noise-driven model were significantly different from the control (*p <* 0.05) for a prolonged time periods. Such differences were not significant under all the other conditions, except for the plastic and spontaneous model in the late epochs, when the model performance under the control condition dropped to the chance level.

**Fig 19.**
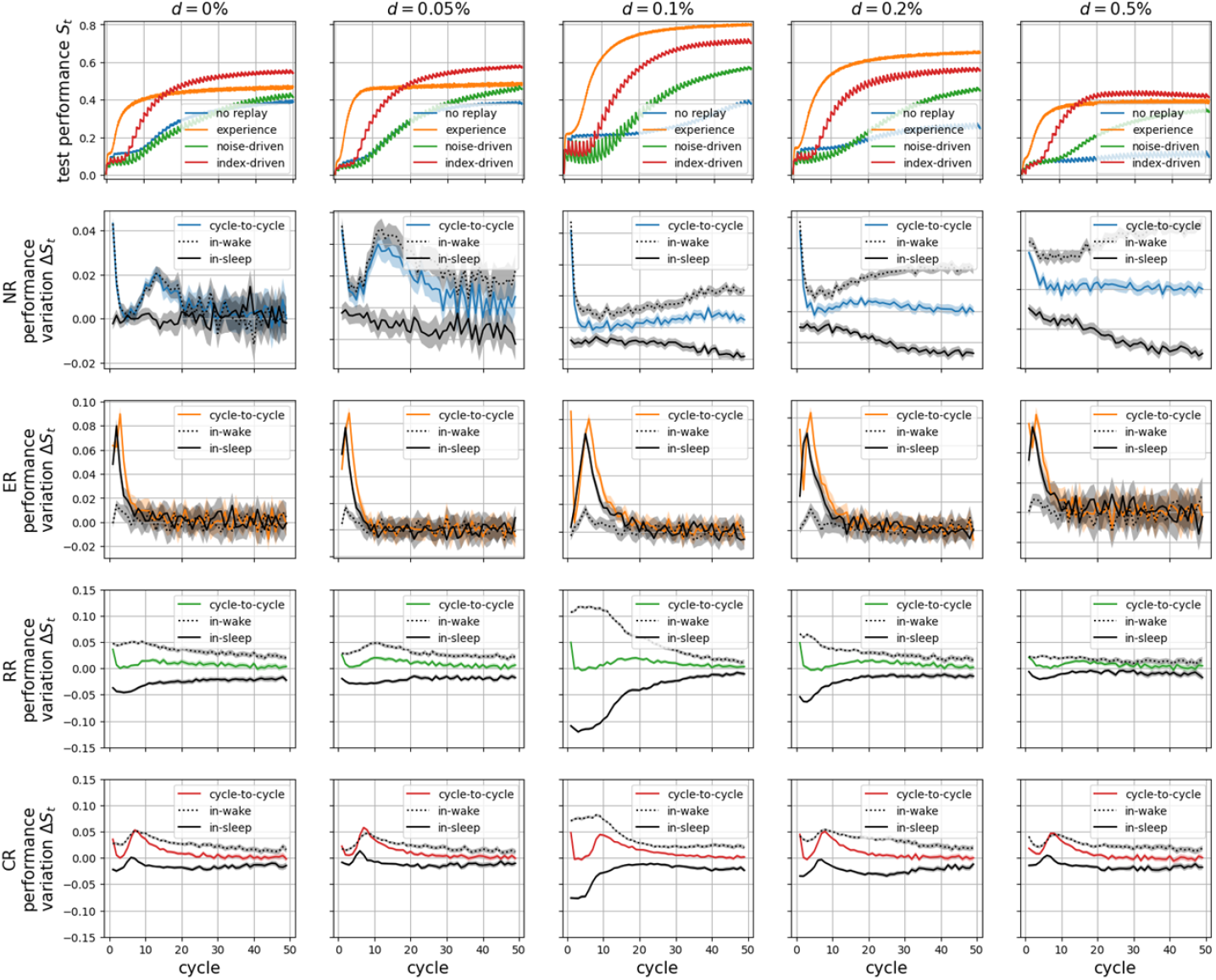
The performance and its cyclic changes of the birdsong development models with different downscaling rates. Although the downscaling rate changed quantitatively the overall performance in all the models, their qualitative behaviours stayed the same. In particular, the index-driven model manifested the most realistic performance oscillation under all the conditions.

## Footnotes

1 The term post-sleep performance deterioration, rather than overnight performance deterioration, is used in [16], presumably because it is impossible to measure performance of real birds in sleep. However, we could, and we did, test our computational models in sleep with all their parameters fixed.

2 We choose the synaptic decay rate no larger than 1% according to [66] but not the L2 regularisation recommended for RBMs. Instead, we use the simple weight decay rule (14) as in [64, 72], which is equivalent to the L2 regularisation in many but not all scenarios [57].

3 It is sensible to use *S*_*t*_ or *S*_*c*_ for the index-driven replay. However, they are not useful for fair comparison to other replay schemes.

## Reference

1. Hayes TL, Krishnan GP, Bazhenov M, Siegelmann HT, Sejnowski TJ, Kanan C. Replay in deep learning: Current approaches and missing biological elements. Neural Computation. 2021;33(11):2908–2950.

2. Roscow EL, Chua R, Costa RP, Jones MW, Lepora N. Learning offline: memory replay in biological and artificial reinforcement learning. Trends in Neurosciences. 2021;.

3. Wittkuhn L, Chien S, Hall-McMaster S, Schuck NW. Replay in minds and machines. Neuroscience & Biobehavioral Reviews. 2021;129:367–388.

4. Pavlides C, Winson J. Influences of hippocampal place cell firing in the awake state on the activity of these cells during subsequent sleep episodes. Journal of neuroscience. 1989;9(8):2907–2918.

5. Qin YL, McNaughton BL, Skaggs WE, Barnes CA. Memory reprocessing in corticocortical and hippocampocortical neuronal ensembles. Philosophical Transactions of the Royal Society of London Series B: Biological Sciences. 1997;352(1360):1525–1533.

6. Louie K, Wilson MA. Temporally structured replay of awake hippocampal ensemble activity during rapid eye movement sleep. Neuron. 2001;29(1):145–156.

7. Lee AK, Wilson MA. Memory of sequential experience in the hippocampus during slow wave sleep. Neuron. 2002;36(6):1183–1194.

8. Ji D, Wilson MA. Coordinated memory replay in the visual cortex and hippocampus during sleep. Nature neuroscience. 2007;10(1):100–107.

9. Tomé DF, Sadeh S, Clopath C. Coordinated hippocampal-thalamic-cortical communication crucial for engram dynamics underneath systems consolidation. Nature Communications. 2022;13(1):840.

10. O’Neill J, Boccara CN, Stella F, Schönenberger P, Csicsvari J. Superficial layers of the medial entorhinal cortex replay independently of the hippocampus. Science. 2017;355(6321):184–188.

11. Székely AD. The avian hippocampal formation: subdivisions and connectivity. Behavioural brain research. 1999;98(2):219–225.

12. Emery NJ, Clayton NS. Evolution of the avian brain and intelligence. Current Biology. 2005;15(23):R946–R950.

13. Striedter GF. Evolution of the hippocampus in reptiles and birds. Journal of Comparative Neurology. 2016;524(3):496–517.

14. Stacho M, Herold C, Rook N, Wagner H, Axer M, Amunts K, et al. A cortex-like canonical circuit in the avian forebrain. Science. 2020;369(6511):eabc5534.

15. Dave AS, Margoliash D. Song replay during sleep and computational rules for sensorimotor vocal learning. Science. 2000;290(5492):812–816.

16. Derégnaucourt S, Mitra PP, Fehér O, Pytte C, Tchernichovski O. How sleep affects the developmental learning of bird song. Nature. 2005;433(7027):710–716.

17. Káli S, Dayan P. Off-line replay maintains declarative memories in a model of hippocampal-neocortical interactions. Nature neuroscience. 2004;7(3):286–294.

18. Squire LR, Genzel L, Wixted JT, Morris RG. Memory consolidation. Cold Spring Harbor perspectives in biology. 2015;7(8):a021766.

19. Buhry L, Azizi AH, Cheng S. Reactivation, replay, and preplay: how it might all fit together. Neural plasticity. 2011;2011.

20. Ólafsdóttir HF, Bush D, Barry C. The role of hippocampal replay in memory and planning. Current Biology. 2018;28(1):R37–R50.

21. McClelland JL, McNaughton BL, O’Reilly RC. Why there are complementary learning systems in the hippocampus and neocortex: insights from the successes and failures of connectionist models of learning and memory. Psychological review. 1995;102(3):419.

22. French RM. Catastrophic forgetting in connectionist networks. Trends in cognitive sciences. 1999;3(4):128–135.

23. Grossberg S. Processing of expected and unexpected events during conditioning and attention: a psychophysiological theory. Psychological review. 1982;89(5):529.

24. Kumaran D, Hassabis D, McClelland JL. What learning systems do intelligent agents need? Complementary learning systems theory updated. Trends in cognitive sciences. 2016;20(7):512–534.

25. McClelland JL, McNaughton BL, Lampinen AK. Integration of new information in memory: new insights from a complementary learning systems perspective. Philosophical Transactions of the Royal Society B. 2020;375(1799):20190637.

26. Lewis PA, Bendor D. How targeted memory reactivation promotes the selective strengthening of memories in sleep. Current Biology. 2019;29(18):R906–R912.

27. Navarrete M, Greco V, Rakowska M, Bellesi M, Lewis PA. Auditory stimulation during REM sleep modulates REM electrophysiology and cognitive performance. Communications Biology. 2024;7(1):193.

28. Shepard RN. Stimulus and response generalization: A stochastic model relating generalization to distance in psychological space. Psychometrika. 1957;22(4):325–345.

29. Nosofsky RM. Attention, similarity, and the identification–categorization relationship. Journal of experimental psychology: General. 1986;115(1):39.

30. Smolensky P. 6. In: Rumelhart DE, McClclland JL, the PDP Research Group, editors. Information processing in dynamical systems: Foundations of harmony theory. MIT Press; 1986. p. 194–281.

31. Hinton GE. Training products of experts by minimizing contrastive divergence. Neural computation. 2002;14(8):1771–1800.

32. Bendor D, Wilson MA. Biasing the content of hippocampal replay during sleep. Nature neuroscience. 2012;15(10):1439–1444.

33. Oudiette D, Antony JW, Creery JD, Paller KA. The role of memory reactivation during wakefulness and sleep in determining which memories endure. Journal of Neuroscience. 2013;33(15):6672–6678.

34. Oudiette D, Paller KA. Upgrading the sleeping brain with targeted memory reactivation. Trends in cognitive sciences. 2013;17(3):142–149.

35. Cairney SA, Lindsay S, Sobczak JM, Paller KA, Gaskell MG. The benefits of targeted memory reactivation for consolidation in sleep are contingent on memory accuracy and direct cue-memory associations. Sleep. 2016;39(5):1139–1150.

36. Schechtman E, Heilberg J, Paller KA. Memory consolidation during sleep involves context reinstatement in humans. Cell reports. 2023;42(4).

37. Rakowska M, Abdellahi ME, Bagrowska P, Navarrete M, Lewis PA. Long term effects of cueing procedural memory reactivation during NREM sleep. Neuroimage. 2021;244:118573.

38. Rakowska M, Bagrowska P, Lazari A, Navarrete M, Abdellahi ME, Johansen-Berg H, et al. Cueing memory reactivation during NREM sleep engenders long-term plasticity in both brain and behaviour. Imaging Neuroscience. 2024;2:1–21.

39. Lerner I, Lupkin SM, Tsai A, Khawaja A, Gluck MA. Sleep to remember, sleep to forget: Rapid eye movement sleep can have inverse effects on recall and generalization of fear memories. Neurobiology of Learning and Memory. 2021;180:107413.

40. Albouy G, Sterpenich V, Balteau E, Vandewalle G, Desseilles M, Dang-Vu T, et al. Both the hippocampus and striatum are involved in consolidation of motor sequence memory. Neuron. 2008;58(2):261–272.

41. Buzsáki G, Tingley D. Space and time: the hippocampus as a sequence generator. Trends in cognitive sciences. 2018;22(10):853–869.

42. Stoianov I, Maisto D, Pezzulo G. The hippocampal formation as a hierarchical generative model supporting generative replay and continual learning. Progress in Neurobiology. 2022;217:102329.

43. Teyler TJ, DiScenna P. The hippocampal memory indexing theory. Behavioral neuroscience. 1986;100(2):147.

44. Robins A. Catastrophic forgetting, rehearsal and pseudorehearsal. Connection Science. 1995;7(2):123–146.

45. Stickgold R. Sleep-dependent memory consolidation. Nature. 2005;437(7063):1272–1278.

46. Diekelmann S, Born J. The memory function of sleep. Nature Reviews Neuroscience. 2010;11(2):114–126.

47. Singh D, Norman KA, Schapiro AC. A model of autonomous interactions between hippocampus and neocortex driving sleep-dependent memory consolidation. Proceedings of the National Academy of Sciences. 2022;119(44):e2123432119.

48. Deperrois N, Petrovici MA, Senn W, Jordan J. Learning cortical representations through perturbed and adversarial dreaming. Elife. 2022;11:e76384.

49. Kudrimoti HS, Barnes CA, McNaughton BL. Reactivation of hippocampal cell assemblies: effects of behavioral state, experience, and EEG dynamics. Journal of Neuroscience. 1999;19(10):4090–4101.

50. van de Ven GM, Siegelmann HT, Tolias AS. Brain-inspired replay for continual learning with artificial neural networks. Nature communications. 2020;11(1):1–14.

51. Mocanu DC, Vega MT, Eaton E, Stone P, Liotta A. Online contrastive divergence with generative replay: Experience replay without storing data. arXiv preprint arXiv:161005555. 2016;.

52. Genzel L, Robertson EM. To replay, perchance to consolidate. PLoS biology. 2015;13(10):e1002285.

53. Yonelinas AP, Ranganath C, Ekstrom AD, Wiltgen BJ. A contextual binding theory of episodic memory: systems consolidation reconsidered. Nature Reviews Neuroscience. 2019;20(6):364–375.

54. Dudai Y. The neurobiology of consolidations, or, how stable is the engram? Annu Rev Psychol. 2004;55:51–86.

55. Parisi GI, Kemker R, Part JL, Kanan C, Wermter S. Continual lifelong learning with neural networks: A review. Neural Networks. 2019;113:54–71.

56. Delange M, Aljundi R, Masana M, Parisot S, Jia X, Leonardis A, et al. A continual learning survey: Defying forgetting in classification tasks. IEEE Transactions on Pattern Analysis and Machine Intelligence. 2021;.

57. Loshchilov I, Hutter F. Decoupled weight decay regularization. arXiv preprint arXiv:171105101. 2017;.

58. Zhang G, Wang C, Xu B, Grosse R. Three mechanisms of weight decay regularization. arXiv preprint arXiv:181012281. 2018;.

59. Liu Z, Sun M, Zhou T, Huang G, Darrell T. Rethinking the value of network pruning. arXiv preprint arXiv:181005270. 2018;.

60. Blalock D, Gonzalez Ortiz JJ, Frankle J, Guttag J. What is the state of neural network pruning? Proceedings of machine learning and systems. 2020;2:129–146.

61. Tononi G, Cirelli C. Sleep and synaptic homeostasis: a hypothesis. Brain research bulletin. 2003;62(2):143–150.

62. Tononi G, Cirelli C. Sleep function and synaptic homeostasis. Sleep medicine reviews. 2006;10(1):49–62.

63. Sullivan TJ, de Sa VR. Sleeping our way to weight normalization and stable learning. Neural computation. 2008;20(12):3111–3130.

64. Hill S, Tononi G, Ghilardi MF. Sleep improves the variability of motor performance. Brain research bulletin. 2008;76(6):605–611.

65. Tononi G, Cirelli C. Sleep and the price of plasticity: from synaptic and cellular homeostasis to memory consolidation and integration. Neuron. 2014;81(1):12–34.

66. Hinton GE. A practical guide to training restricted Boltzmann machines. In: Neural networks: Tricks of the trade. Springer; 2012. p. 599–619.

67. Marr D. Simple memory: a theory for archicortex. Philosophical Transactions of the Royal Society of London B, Biological Sciences. 1971;262(841):23–81. doi:10.1098/rstb.1971.0078.

68. Willshaw D, Dayan P, Morris RG. Memory, modelling and Marr: a commentary on Marr (1971)’Simple memory: a theory of archicortex’. Philosophical Transactions of the Royal Society B: Biological Sciences. 2015;370(1666):20140383.

69. Kumaran D, McClelland JL. Generalization through the recurrent interaction of episodic memories: a model of the hippocampal system. Psychological review. 2012;119(3):573.

70. Luce RD. The choice axiom after twenty years. Journal of mathematical psychology. 1977;15(3):215–233.

71. Luce RD. Individual choice behavior: A theoretical analysis. Courier Corporation; 2012.

72. Krogh A, Hertz J. A simple weight decay can improve generalization. Advances in neural information processing systems. 1991;4.

73. Choi SS, Cha SH, Tappert CC. A survey of binary similarity and distance measures. Journal of systemics, cybernetics and informatics. 2010;8(1):43–48.

74. Brusco M, Cradit JD, Steinley D. A comparison of 71 binary similarity coefficients: The effect of base rates. Plos one. 2021;16(4):e0247751.

75. Lucic M, Kurach K, Michalski M, Gelly S, Bousquet O. Are gans created equal? a large-scale study. Advances in neural information processing systems. 2018;31.

76. Sajjadi MS, Bachem O, Lucic M, Bousquet O, Gelly S. Assessing generative models via precision and recall. Advances in Neural Information Processing Systems. 2018;31.

77. Pielou EC. The measurement of diversity in different types of biological collections. Journal of theoretical biology. 1966;13:131–144.

